# Leveraging AI and structural proteomics for rational design of a KAT6A degrader

**DOI:** 10.64898/2026.05.25.727609

**Authors:** Gali Arad, Nitzan Simchi, Sagie Brodsky, Alon Shtrikman, Yonatan Kedem, Iris Alchanati, Galina Otonin, Anjana Shenoy, Dimitri Kovalerchik, Michal Ran Shchory, Yaron Ben Shoshan-Galeczki, Noam Cohen, Katharina Lange, Eran Seger, Kirill Pevzner

## Abstract

While targeted protein degraders such as PROTACs are a clinically proven therapeutic strategy, the discovery of novel degraders remains hampered by trial-and-error process. To address this challenge, we developed the AIMS™ platform, which combines structural proteomics with AI models for rational PROTAC design. AIMS™ is an end-to-end toolkit for PROTAC optimization, encompassing structure solving using proteomics and AI, prediction of ADME and degradation properties, and prospective ranking of compound design ideas. Altogether, this integrated platform successfully enabled the multi-parameter optimization of a potent and bioavailable *in vivo* validated KAT6A degrader, establishing a versatile framework for PROTAC development across various targets.

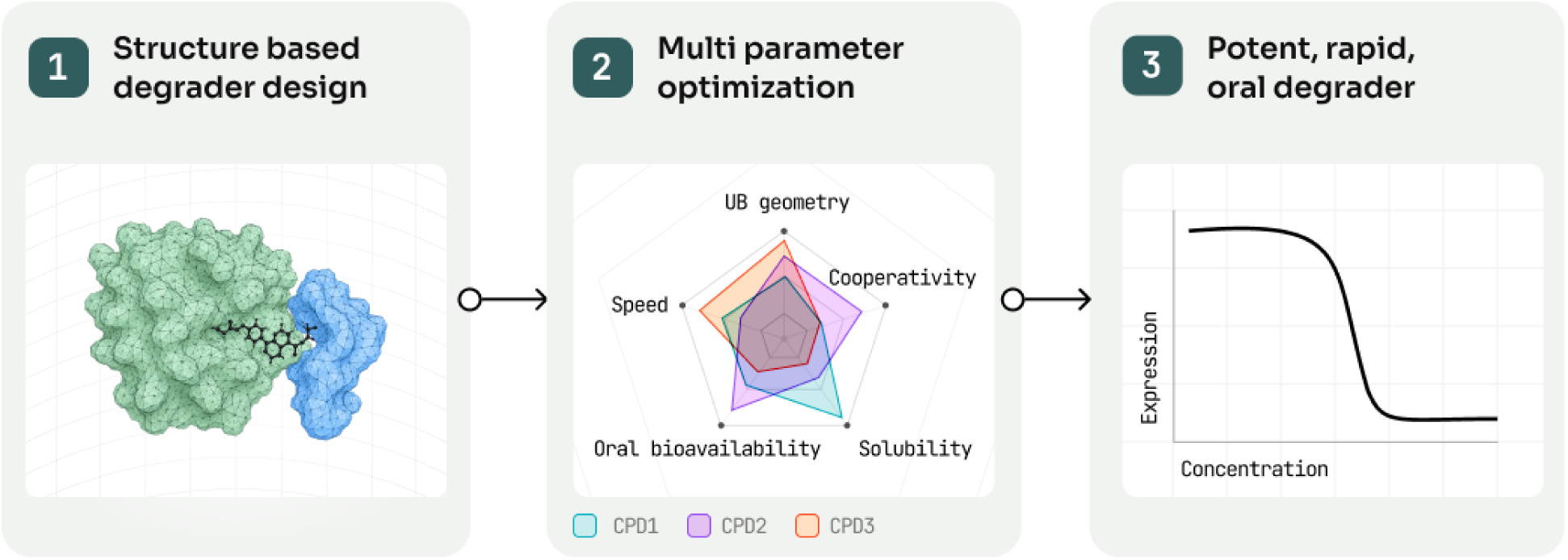

## Introduction

Proximity-inducing drugs are an emerging class of protein interaction modulators, most famously exemplified by proteolysis-targeting chimera (PROTAC). PROTACs are heterobifunctional compounds consisting of two ligands joined by a linker to bring together an E3 ligase and a target protein of interest (POI) to induce ubiquitination and subsequent degradation of the latter^1,2^. Alongside the estrogen receptor (ER) degrader ARV-471 -which recently became the first PROTAC to receive FDA approval^3^ -dozens of other PROTAC drugs are currently being investigated in clinical trials, including the androgen receptor (AR) degrader BMS-986365^4^ and others^5^. While promising, the complexity of PROTAC optimization has proven challenging and has led researchers to rely mostly on trial and error, thus slowing down their translation into clinical applications^6–7^.

Conventional approaches hinder PROTAC development since the activity of a PROTAC molecule is driven by multiple factors. While efficacy of classical small molecule inhibitors is often mediated by the binding affinity of a single molecule to a single protein, PROTAC efficacy is driven by the assembly and the dynamics of a POI-E3-PROTAC ternary complex^8^. The ternary complex dynamics require a balance between complex stability and flexibility to enable efficient degradation. This is further complicated by protein-protein interactions formed within the ternary complex, leading to some compounds inducing more cooperative complexes than others; which is, however, not consistently correlated with improved degradation activity^8–10^. The ubiquitination geometry of the complex, namely how well the recipient lysine residue is located relative to the E3 ligase complex, is another mechanism affecting degradation, supported by emerging experimental evidence^11,12^. On top of these, oral bioavailability and permeability represent major challenges in PROTAC development, due to the heterobifunctional structure and high molecular weight of PROTACs, which places them outside the “Rule of 5” and leading to poor solubility and low permeability^13–15^. It is therefore clear that degradation efficiency is a multi-faceted challenge that cannot be determined by any single factor alone, highlighting the need for the development of structure-based, computational drug design methodologies.

For rational PROTAC design, deep understanding of the conformational dynamics of complex assembly is required^16,17^, and it is likely that several conformations represent valid structures, rather than a single static one. Experimental approaches such as X-ray crystallography or Cryogenic electron microscopy (Cryo-EM) provide important structural information required for PROTAC design, as shown by Gadd and colleagues in the case of BRD4 PROTAC^18^ or by Nayak and colleagues in the case of BCL-xL/BCL-2^18,19^. Despite these compelling examples, these methods are static by nature, therefore ignoring ternary complex dynamics. Specifically, cryo-EM can offer a glimpse into complex dynamics by capturing different states, but this still represents a fragmented view ignoring the continuous conformational landscape. Moreover, these approaches are slow, low-throughput, and costly, limiting their compatibility with accelerated drug development workflows and their utility in supporting structure–activity relationship (SAR) studies^20^.

Modeling ternary complexes has also been addressed by computational methods, including protein-protein docking, physics-based tools, and generative AI models^21^. Molecular docking is a popular technique used for structure based drug design, but its application for ternary complex modeling is hindered by insufficient sampling and failure to accurately represent flexible and dynamic complex behavior. Molecular dynamics (MD) simulations are used to assess ternary complex stability and refine potential structures. While this allows some level of conformational dynamics, these simulations are inherently limited by their structural starting point, as well as simulation duration, parameterization, and high computational costs^22,23^. AI models, most notably AlphaFold3 and Boltz2^24–26^, provide essentially proteome-wide structure predictions. However, protein complexes and interactions have traditionally been hard to model as they are sparsely represented in existing protein structure resources (e.g. Protein Data Bank [PDB], used as one of the main sources for model training). Thus, while demonstrating overall high confidence across the entire proteome, they also possess the risk of overconfidence in prediction of a given protein or complex, for example in predictions of antibody-antigen complexes^27^ or in cases of multiple conformations^28,29^, and particularly in the case of ternary complex prediction, where these models have been repeatedly shown to underperform^30,31^.

Structural proteomics methods such as cross-linking mass spectrometry (XL-MS) and hydrogen–deuterium exchange mass spectrometry (HDX-MS) enable the analysis of proteins and complexes in their native state^32–34^, thus overcoming limitations of other experimental methods and providing access to dynamic conformational ensembles. Indeed, there is emerging evidence of the implementation of structural MS methodologies in drug discovery workflows, either in hit discovery enablement^35,36^ or in later stages of drug optimization^37,38^. However, this evidence represents small scale examples, and structural MS techniques have yet to be applied to drug discovery systematically. These limitations collectively demonstrate that a rational approach to PROTAC design cannot rely solely on either experimental or AI-based methods. Instead, optimal PROTAC design requires leveraging the speed and scalability of AI models in combination with experimental data.

Beyond the challenges of structural modeling, additional computational efforts have focused on optimizing critical pharmacological parameters such as absorption, distribution, metabolism, and excretion (ADME) and degradation kinetics. Machine learning-based models have been developed to predict cell permeability and oral bioavailability, often utilizing molecular descriptors such as lipophilicity and topological polar surface area^39^. Other tools, like PROTAC-TS, employ generative chemical language models combined with reinforcement learning to design linkers that specifically improve membrane permeability^40^. Furthermore, mechanistic pharmacodynamic frameworks have emerged to model the “hook effect” and quantify the relationship between target occupancy and degradation speed^41^. However, while these individual tools address specific hurdles in the PROTAC development pipeline, they are not designed as an integrated, end-to-end suite of tools capable of optimizing the complex interplay between structural dynamics, catalytic efficiency, and drug-like properties.

In this work, we introduce the AIMS™ platform, where we leverage structural proteomics combined with proprietary AI models and physics-based simulations, and show its application in the development and optimization of a KAT6A degrader, a biologically and clinically validated target in breast cancer and other malignancies^42^. AIMS™ platform consists of multiple modules that are uniquely designed for particular PROTAC optimization tasks such as binding affinity, degradation speed, and bioavailability. This modular approach enabled a multi-parameter optimization strategy, resulting in the accelerated transformation of an *in vitro* compound into a potent, orally bioavailable KAT6A degrader. To our knowledge, this represents the first detailed report of structure-based degrader design in the context of a commercial drug discovery project, providing a blueprint for the future of proximity-modulating therapeutics including PROTACs, molecular glues, and others.

## Results

### AIMS-Fold: PROTAC-dependent structural features identify a productive conformation

Degrader activity largely depends on the proper assembly of a ternary complex, composed of the protein of interest (POI), an E3 ligase (e.g. most commonly CRBN, VHL) and a heterobifunctional compound inducing their proximity. Solving the KAT6A–PROTAC–CRBN ternary complex structure is central to rational design, as it provides the structural blueprint necessary to distinguish productive degradation assemblies from unproductive ones.

To study the ternary complex structure of KAT6A degraders, we took an initial set of compounds with different levels of activity, defined by degradation efficiency as either strong (DC50 < 50 nM) or weak (DC50 > 1000 nM). As those PROTACs differ by the linker length, flexibility and attachment points to the warheads, we hypothesized that strong degraders will lead to ternary complex conformations that are different from the ones formed by weaker ones. To test this, we applied both cross-linking mass spectrometry (XL-MS) and hydrogen deuterium exchange (HDX-MS) to study the KAT6A-CRBN-compound complex. Indeed, both MS datasets identified PROTAC-dependent differences between strong and weak degraders. The XL-MS data yielded specific inter-protein crosslinks that are unique to strong degraders (**Figure 1A and Supplementary Figure S1A-B**), suggesting that a specific KAT6A orientation relative to CRBN is required for a productive ubiquitination to occur. Furthermore, HDX-MS data identified regions with differential levels of surface exposure in accordance with compound activity (**Figure 1B and Supplementary Figure S2**). Specifically, region A is more protected in strong compounds, while region B is more protected in weak compounds.

**Figure 1.**
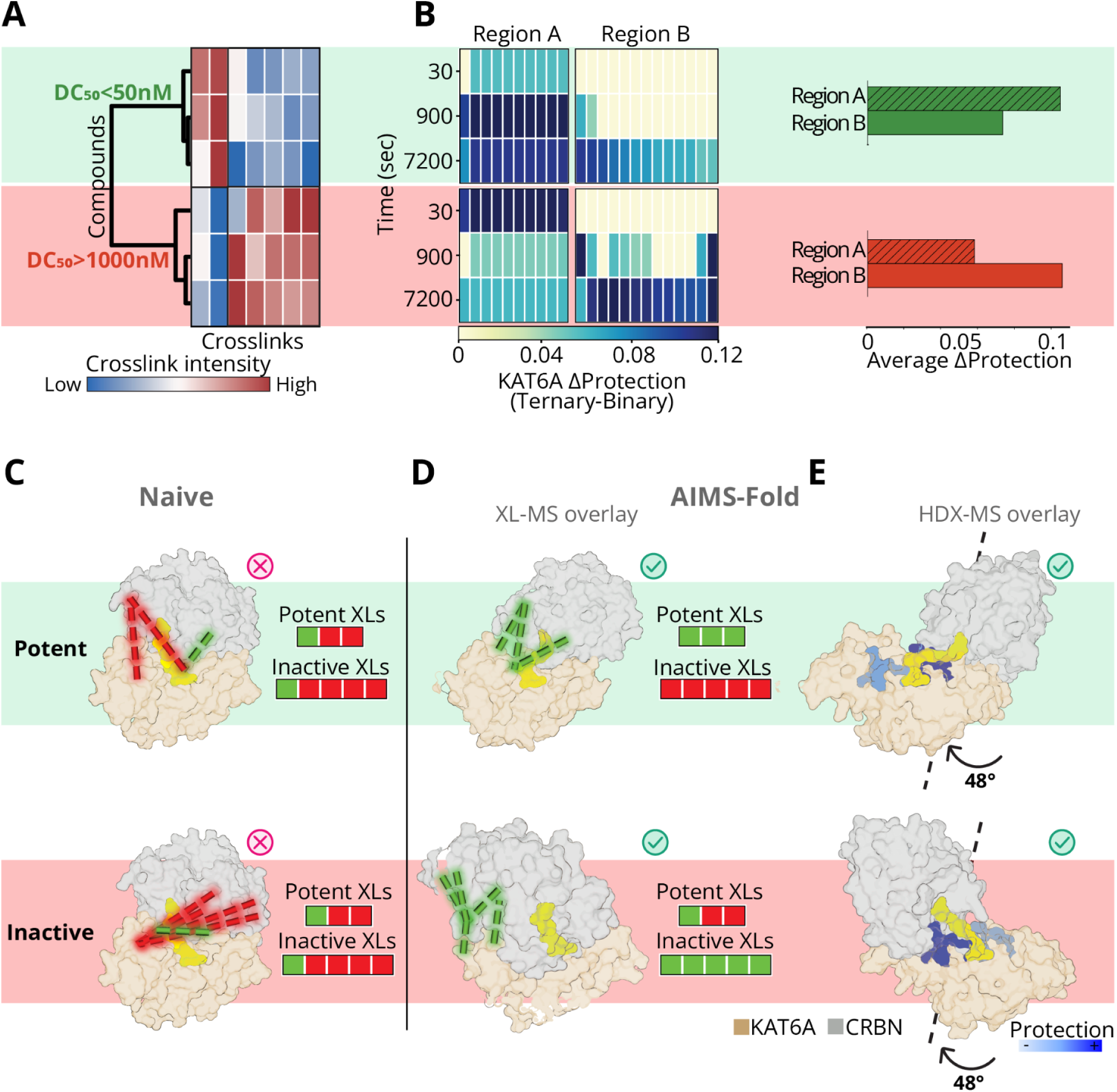
AIMS™ reveals conformational differences between strong and weak degraders. A- Heatmap of XL-MS results describing crosslinks significantly differentiating weak and strong compounds. Different columns represent different crosslinks and color represents crosslink signal intensity. B- Heatmap of HDX-MS results describing the protection pattern of two KAT6A regions, comparing strong (top) and weak (bottom) compounds. Each box represents a single residue. Darker colors indicate stronger protection, namely KAT6A regions that are buried upon ternary complex formation. The barplot on the right describes the signal per region in the latest timepoint. C-D- Boltz-2 prediction of KAT6A-CRBN ternary complex with potent (top) or weak (bottom) degraders, without (C) or with (D) MS-derived constraints. Dashed lines represent crosslinks identified by XL-MS data, and color indicates whether they were satisfied by the model (green) or violated (red). Each conformation displays only its state specific crosslinks (three for the strong models and five for the weak). Notably, one of the crosslinks defining potent compounds was imputed from KAT6B (Supplementary Figure S1B-D). E- HDX-MS derived protected areas are overlaid on ternary complex conformations of strong and weak degraders (different angle of conformations shown in D, as indicated by the rotation arrow), different regions protected in each conformation are marked in color. Darker blue shades indicate stronger protection. Dark blue region in top structure indicates Region A, while dark blue region in bottom structure indicates Region B.

To identify the productive ternary complex conformation for KAT6A degradation, we used AIMS-Fold, where structural MS studies are coupled with AI modeling to identify ternary structures induced by multiple compounds^43^. Briefly, AIMS-Fold is a diffusion based generative model for biomolecular structure prediction with extended support for structural proteomics constraints. Translating the MS results into structural constraints enabled informed structural predictions of the various ternary complexes. Notably, while naïve AI models violate the majority of constraints regardless of compound (**Figure 1C**), AIMS-Fold model identifies a productive conformation and a non-productive conformation -satisfying both the XL-MS constraints as well as the HDX derived protection patterns (**Figure 1D,E**). Unlike existing structural AI (Boltz-2, AlphaFold3 etc.) models, AIMS-Fold takes into account both “positive” and “negative” distance constraints -namely residues that should be in close proximity (attracted) and residues that should be kept apart (repelled). Importantly, not only does the productive conformation satisfy the crosslinks associated with strong degraders, it also does *not* satisfy the ones associated with weaker ones, made possible due to the repel constraints feature of AIMS-Fold (see Methods).

Ternary complexes are often not represented by a single static state. Therefore, conformational dynamics is also expected to play a role in degradation efficiency. Remarkably, this notion was supported by the HDX-MS time course data showing that differentiating protection patterns increase over time (**Figure 1B**). To evaluate the stability of the productive conformation, we explored the conformational free energy landscape using metadynamics simulations. We defined our collective variables (CVs) based on key interface distances and torsional angles (see Methods). The resulting probability distributions in the CV space allowed us to distinguish between free energy surfaces for the evaluated degraders, characterized by two primary conformational basins in the upper left quadrant (Q1) and in the bottom right quadrant (Q4, **Figure 2A and Supplementary Video 1**). Interestingly, we observed a clear association between basin occupancy and degrader efficacy; occupancy in the Q4 region of the CV space is lower for a strong degrader (**Figure 2A, top**), while increased occupancy in the Q4 region is characteristic of a weak degrader (**Figure 2A, bottom**). Taken together, AIMS-Fold discovered a productive KAT6A PROTAC conformation required for potent degradation. Metadynamics simulations further supported this finding and showed that the tendency to maintain a certain conformation reflects the activity level of the compound.

**Figure 2.**
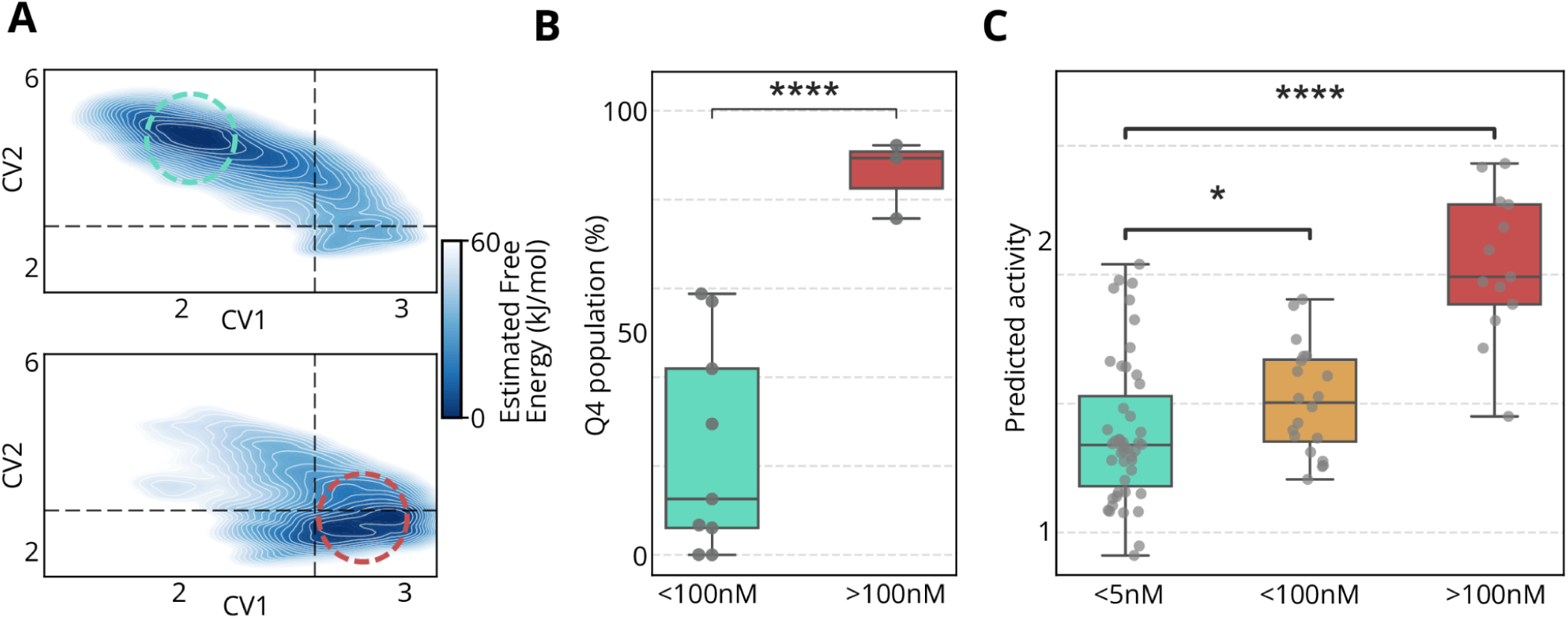
AIMS-Rank translates structural insight into a high throughput virtual screening capability. A- Differential estimated free energy landscapes indicate conformational dynamics for different compounds, and show stabilization of the active compound in the upper left quadrant (top, basin marked in green dashed circle); and the inactive compound in the lower right quadrant (bottom, basin marked in red dashed circle). See also Supplementary Video 1. B- Boxplot summarizing multiple metadynamics simulations of multiple compounds, indicating the association between productive state (measured by occupancy of Q4 in the estimated free energy landscape) and degradation efficiency. C- Association between measured and predicted activity, enabled by AIMS™ structures. Colors indicate weak (red), slightly active (yellow) and active (green) degraders. Lower y-axis values indicate better predicted activity.

### AIMS-Rank: Virtual screening based on productive conformations enables compound prioritization

We next sought to develop a computational tool allowing us to leverage our diverse structural models to predict degrader activity. Such a tool, enabling prioritization of compounds, would significantly reduce the need for trial and error synthesis and laborious experimental evaluation of large numbers of compounds. The productive conformation we identified, supported by the difference in estimated free energy landscapes (**Figure 1**), encouraged us to use our models in a predictive manner. To further validate the association between productive conformation and degrader activity, we ran multiple metadynamics simulations with additional compounds, confirming that the productive state explains degradation efficacy (**Figure 2B and Supplementary Figure S3**).

While highly informative, metadynamics simulations are computationally expensive and lack the throughput required for large scale virtual screening. In order to overcome this limitation and generate an efficient compound ranking model, we utilized the features describing productive and unproductive conformations as derived from our AIMS-Fold generated models and dynamic simulations, and developed a co-folding based affinity prediction approach. This approach integrates the PROTAC-dependent conformations into our predictive models. We used AIMS-Fold together with the Boltz-2 affinity module to rank and prioritize compounds for synthesis. For each compound, we generated multiple ternary prediction models and calculated the predicted affinity only for those that resemble the active conformation, which is the conformation we wish to stabilize (see Methods). Comparing the predicted activity scores against experimental activity, we found that our model accurately separates active compounds from inactive ones (Pearson *r* = 0.58; **Figure 2C**). The model also significantly differentiates highly active from moderately active compounds, supporting faster early lead optimization.

Overall, these results demonstrate that the differentiation between productive and unproductive states provides a robust framework for high-throughput ranking, showcasing the ability of AIMS-Rank to translate deep structural insights from a limited dataset into a virtual screening capability.

### Active conformations lead to favorable ubiquitination geometry

Having identified a productive conformation, we next wanted to explore the underlying cause contributing to its productivity. One such cause might be the increased time Ub-receptive lysines reside in proximity to the E2-Ub complex, as a higher target degradation rate is fundamentally linked to increased ubiquitination or an increased number of ubiquitination sites accessible to the proteasomal degradation machinery^44^. To provide initial validation and identify the target lysines of KAT6A, we performed a mass spectrometry ubiquitination enrichment experiment utilizing several compounds with varying degradation potencies and correlated their efficacy (DC50) against total ubiquitin intensity (see Methods). This analysis identified several lysine residues significantly ubiquitinated upon treatment, establishing a strong association between total Ub signal and DC50 (**Figure 3A**). We next wanted to test whether target lysines, showing the highest ubiquitination signal following a treatment with a strong PROTAC, are found in high proximity to the E2-Ub complex.

**Figure 3.**
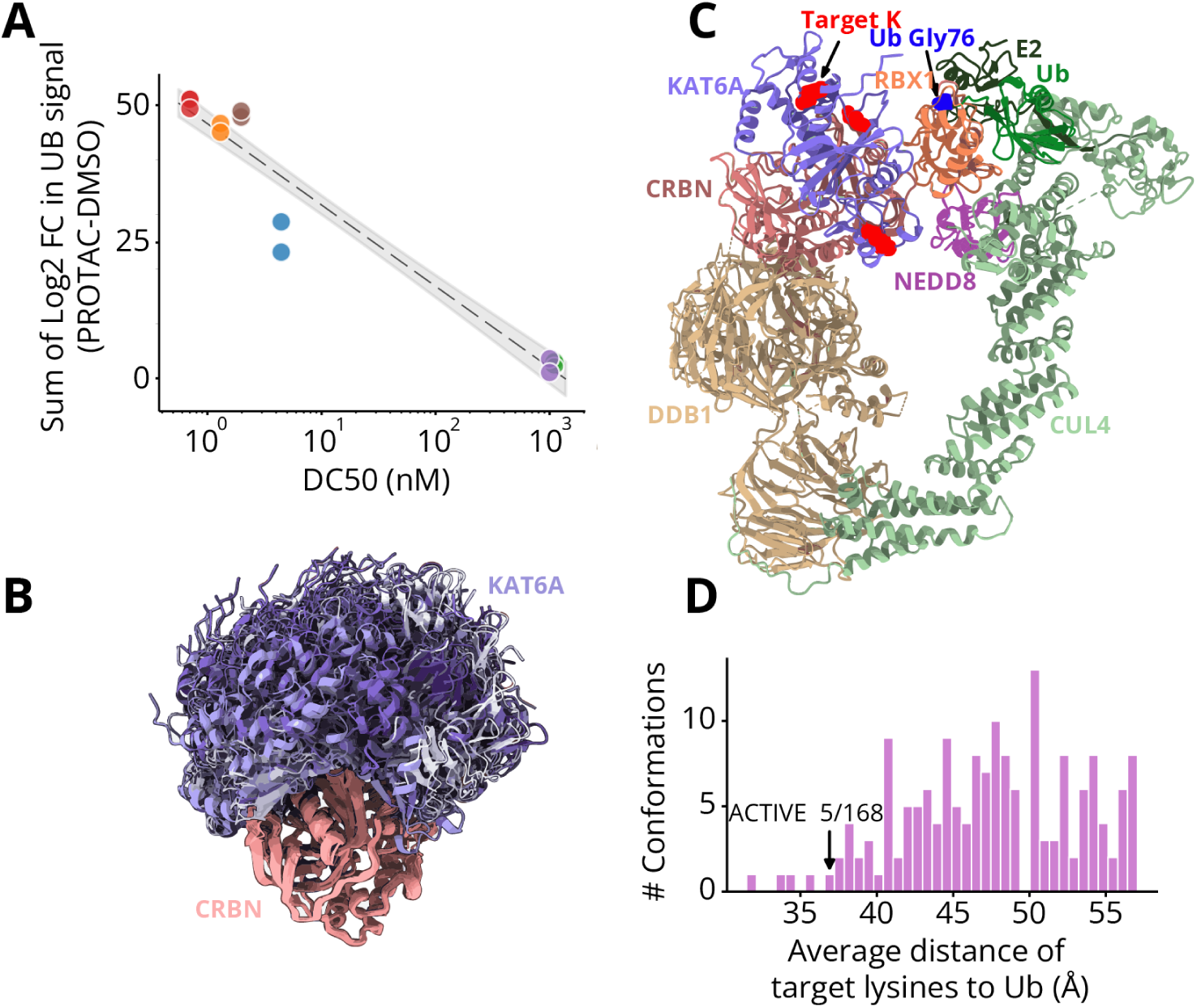
Active conformations lead to favorable ubiquitination geometry. A- Correlation between compound activity (DC50) and the sum of the log2 fold change in Ub intensity between PROTAC treated cells and DMSO. 2 Ub repeats are shown for each PROTAC. B- Ensemble of potential KAT6A-CRBN conformations. Models are aligned to CRBN and different KAT6A conformations are colored in shades of purple. C- KAT6A-CRBN ternary complex overlaid with a full E2-Ub model. KAT6A target lysines are highlighted in red, and Ub Glycine is highlighted in blue. D- histogram of the average distance of the three target lysines from ubiquitin (Gly76) of each ternary conformation in our ensemble. Black arrow points to the AIMS-derived active conformation, ranked 5th overall.

To that end, we first had to overlay our productive and unproductive predicted conformations with the entire ligase complex. Since our models do not include the full ligase machinery, we followed Bai et. al^11^, and constructed 9 models of the full E2-Ub complex (both full ring-forming and open models, see Bai et. al and Methods). Then, to cover the theoretical space of KAT6A-CRBN binding beyond our predicted active and inactive conformations, we generated an ensemble of 184 ternary conformations (**Figure 3B**). This ensemble was generated using Boltz2, Schrödinger’s degrader-centric and protein-centric docking, and pooled with our original AIMS-Fold models, thus allowing us to evaluate the lysine-Ub distance within a broader context. Sixteen conformations were excluded because they showed clashes between the POI and the E2-Ub complex across all 9 models. Interestingly, one of these was our unproductive conformation. The remaining 168 ternary complexes had at least one valid, non clashing overlaid model. Notably, our AIMS-Fold derived productive conformation introduced clashes in only 2/9 models, and showed that the most highly ubiquitinated lysines are indeed facing the E2-Ub in a favorable orientation (**Figure 3C and Supplementary Table S1**).

For each of the valid 168 ternary conformations, we calculated the minimum distance between each target lysine and Ub across its non-clashing models; specifically, between the epsilon-nitrogen of the target lysines and the C-terminal carbon of ubiquitin (Gly76). The average of these values was used to rank the conformations, where smaller distance indicates higher probability of productive ubiquitination. Using this metric, our AIMS-Fold predicted productive conformation was ranked 5th out of the total 168 (**Figure 3D**). Notably, none of the top 4 conformations align with the structural MS input constraints, i.e. not experimentally supported. This relative ranking demonstrates that the AIMS-Fold productive conformation positions the target lysines closer to Ub than almost all other sampled geometries; thus suggesting that the productive conformation can indeed be explained by favorable ubiquitination geometry.

### Speed of ternary complex formation is driven by exit vector topology

While the AIMS-Rank model provides a robust framework for prioritizing compounds, further improvements can be made by considering additional parameters that impact efficacy. Indeed, we identified specific cases where structural predictions did not align with experimental results. For example, one of our active compounds was ranked poorly by the AIMS-Rank model due to a seemingly sub-optimal ternary interface, yet it exhibited a remarkably high association rate (k_a_ = 6.36×10^7^ 1/Ms), potentially compensating for lower stability (**Supplementary Figure S4**).

Following this example, we hypothesized that the speed of complex formation is a central feature of KAT6A degradation activity. A potent degrader needs to induce a productive complex, but its potency also depends on how rapidly it forms, as ternary complex stability is often linked with degradation efficiency^8^. To study the structural determinants of complex formation kinetics computationally, we developed an enhanced sampling approach, to simulate the pre-organization behavior of the initial binary complex (**Figure 4A**). We specifically focused our simulations on the KAT6A-binding end, because our KAT6A warhead has significantly stronger affinity for the degrader than the CRBN one. Using metadynamics simulations, we found that once bound to KAT6A, strong degraders are oriented outwards from the KAT6A surface while weak degraders spend most of the time adhering to the surface of KAT6A. These results suggest that the outward orientation is priming KAT6A to form a complex with CRBN, while the latter potentially hinders ternary complex formation (**Figure 4B**). Estimated free energy landscapes of different compounds show that the conformations of more active compounds mostly have a larger exit vector angle and smaller buried surface area of the linker (i.e. occupy the lower right quadrant, **Figure 4B and Supplementary Figure S5**).

**Figure 4.**
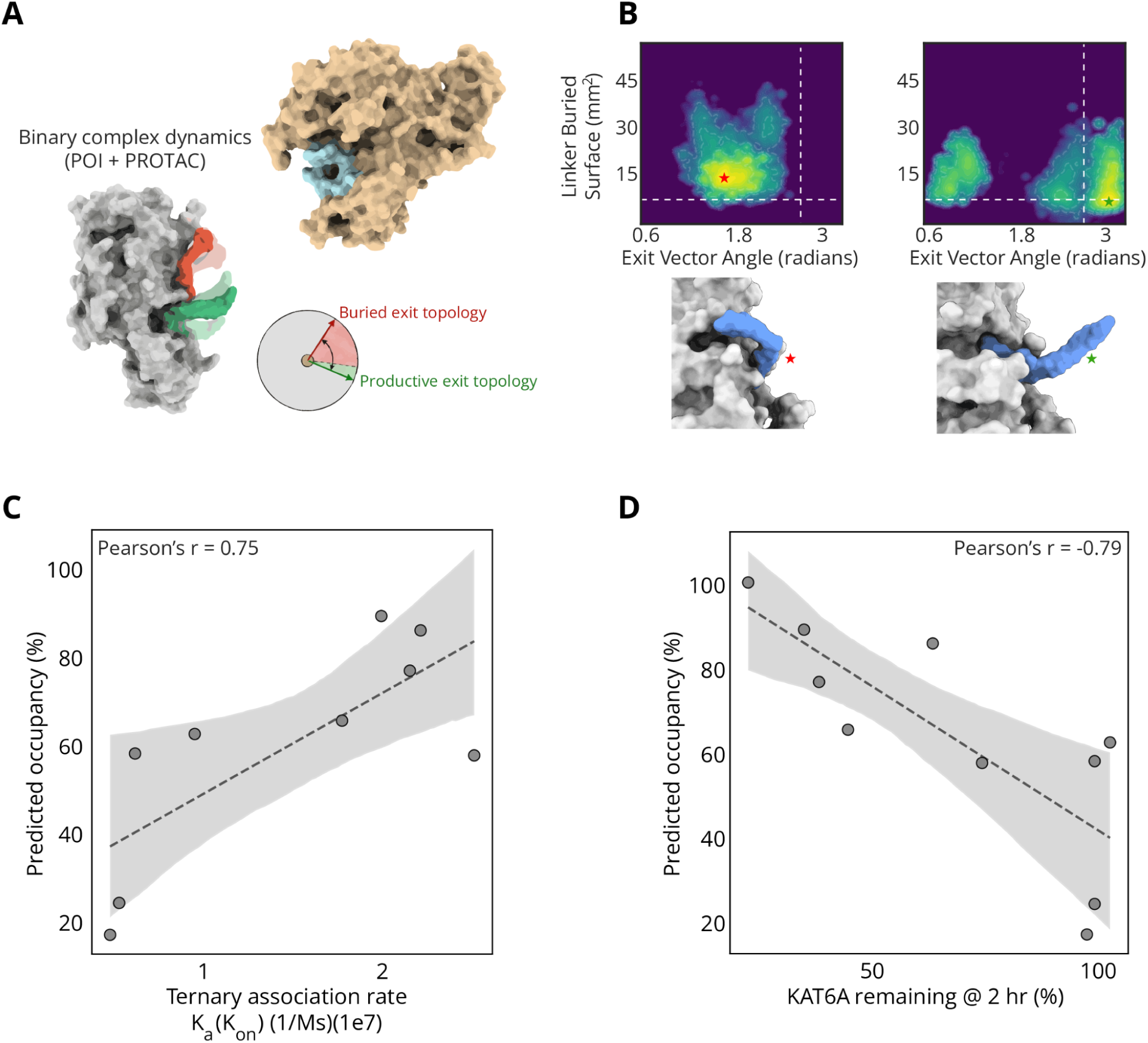
Efficient complex formation is determined by exit vector angle. A- Binary complex dynamics of KAT6A and different PROTACs reveals PROTAC exit vector topology. B- Estimated Free energy landscapes of weak (left) and strong (right) KAT6A degraders, with 3D structures exemplifying exit vector states from each basin noted by a red/ green star. C- Correlation between the productive vector occupancy rate and the ternary complex association rate (K_a_) as measured by SPR. D- Correlation between the productive vector occupancy rate and KAT6A degradation at 2 h (% of Control).

Next, we wanted to test whether these exit topology features can be used to accurately predict PROTAC activity and complex formation. To that end, we initially used compounds for which we had existing experimental data. We calculated their productive exit vector occupancy rate, and found it was correlated (Pearson *r* = 0.75, **Figure 4C**) with the experimental rate of complex association (k_a_ or on-rate, measured by SPR, see Methods). In addition, we used the HiBiT system^45^ on the T47D breast cancer cell line to measure KAT6A levels. To measure degradation speed, we calculated the percentage of KAT6A 2 h post treatment. Consistent with the k_a_ correlation, remaining KAT6A levels were also found to be highly correlated with exit vector topology (Pearson *r* = -0.79, **Figure 4D**), indicating that our model explains both affinity and activity.

### Conformational sampling enables membrane permeability and bioavailability optimization

Having mapped the structural features that guide complex formation and ubiquitination geometry, we next evaluated the cellular bioavailability of additional PROTAC candidates. Improving PROTAC efficacy goes beyond enhancing the PROTAC target affinity or its ability to generate a productive ubiquitination geometry, and achieving sufficient oral bioavailability remains a significant challenge for PROTACs due to their inherent “beyond Rule-of-5” properties. To optimize bioavailability, we evaluated several *in silico* predictors^46^, including features such as logD and membrane permeability^46,47^. This identified a logD prediction model that yielded a strong correlation with experimental data (Pearson *r* = 0.79; **Figure 5A**), underscoring the important role of lipophilicity in driving the bioavailability of our PROTACs.

**Figure 5.**
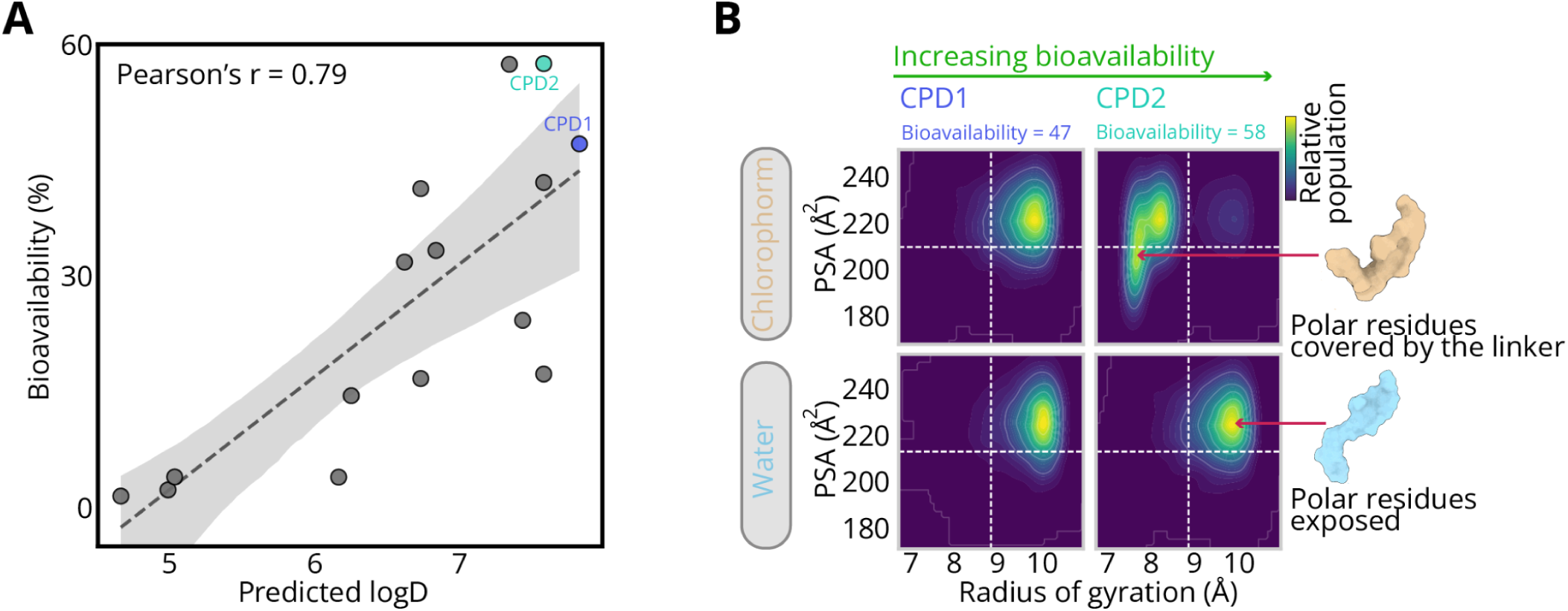
Conformational sampling predicts orally bioavailable compounds. A- Correlation between predicted LogD and experimental bioavailability B- Estimated free energy landscapes demonstrating a reduction in polar surface area in chloroform relative to water for the highly bioavailable PROTAC 2, compared to the less bioavailable PROTAC 1

While the overall correlation between predicted logD and bioavailability was good, we identified several cases where predicted logD does not fully explain bioavailability. One example is CPD1 and CPD2 (**Figure 5A**), where both have similar predicted LogD but different bioavailability. This prompted us to evaluate additional features other than logD. One such feature is chameleonicity, namely the ability to change conformation upon different environmental conditions^15^. To evaluate chameleonic behavior, we used molecular dynamics simulations and modeled the conformational ensembles of our PROTACs across different environments^48^. Specifically, we simulated molecules with varying bioavailability in two distinct solvents: water and chloroform. Water was used to mimic the highly polar aqueous physiological environment of the cell. Chloroform on the other hand, was used to mimic the apolar, lipophilic environment of the cell membrane. To computationally quantify this chameleonic shift, we extracted two key structural descriptors from the ensembles of both solvents: the Polar Surface Area (PSA) and the radius of gyration (Rg). These metrics served as a proxy for the molecule’s ability to shield its polar residues. By calculating the Boltzmann-weighted conformational ensembles in both solvents, we could directly observe how the PROTACs adapt to their environment, namely exposing polar surface areas in an aqueous environment and adopting collapsed, intramolecular hydrogen-bonded conformations in an apolar one. Specifically, the more bioavailable PROTAC (CPD2) was found to modify PSA significantly between the aqueous and chloroform models suggesting a high level of conformation flexibility, i.e. chameleonicity, supported by the linker modification between the two compounds. This effect was not seen with the lower bioavailability PROTAC (CPD1) which maintained similar PSA in both media (**Figure 5B and Supplementary Figure S6**).

### CPD11 is a potent and rapid KAT6A degrader *in vitro* and *in vivo*

We applied our models to facilitate the design of a degrader targeting KAT6A, a biologically and clinically validated target known to play a role in hormone positive (HR+) breast cancer and other malignancies. Rather than yielding a single solution, the AIMS™ platform identified several promising leads that satisfied the diverse requirements of stability, speed, and drug-likeness. Among these, CPD11 was selected for further evaluation as it demonstrated an optimal balance between high predicted ternary stability, efficient kinetic priming, and favorable bioavailability (**Supplementary Figure S7**). We subsequently characterized CPD11 in a series of *in vitro* and *in vivo* assays to validate these platform-derived predictions.

To evaluate the *in vitro* activity of CPD11, T47D breast cancer cell lines with HiBiT-tagged KAT6A were treated for 6 h with CPD11 or vehicle, followed by measurement of intracellular KAT6A levels (see Methods). CPD11 showed excellent degradation efficiency in a dose response assay (DC50 = 0.7 nM) as well as high maximal degradation (Dmax = 89%, **Figure 6A**). In addition, a HiBiT-based kinetics assay where KAT6A levels were measured in different time points throughout a 24 h treatment period, showed KAT6A levels are reduced to 50% during the first 30 minutes of treatment (T_1/2_ = 0.62 h, **Figure 6B**). These results confirmed that CPD11 is a rapid and potent KAT6A degrader. To confirm this was driven by active proteasomal degradation rather than target inhibition alone, we compared KAT6A levels across three conditions: CPD11 alone, CPD11 in combination with the proteasome inhibitor MG132, and CPD11 in combination with an excess of its own KAT6A warhead. KAT6A degradation was significantly higher with CPD11 treatment alone, mechanistically confirming that reduction of KAT6A is proteasome-dependent (**Supplementary Figure S8A**).

**Figure 6.**
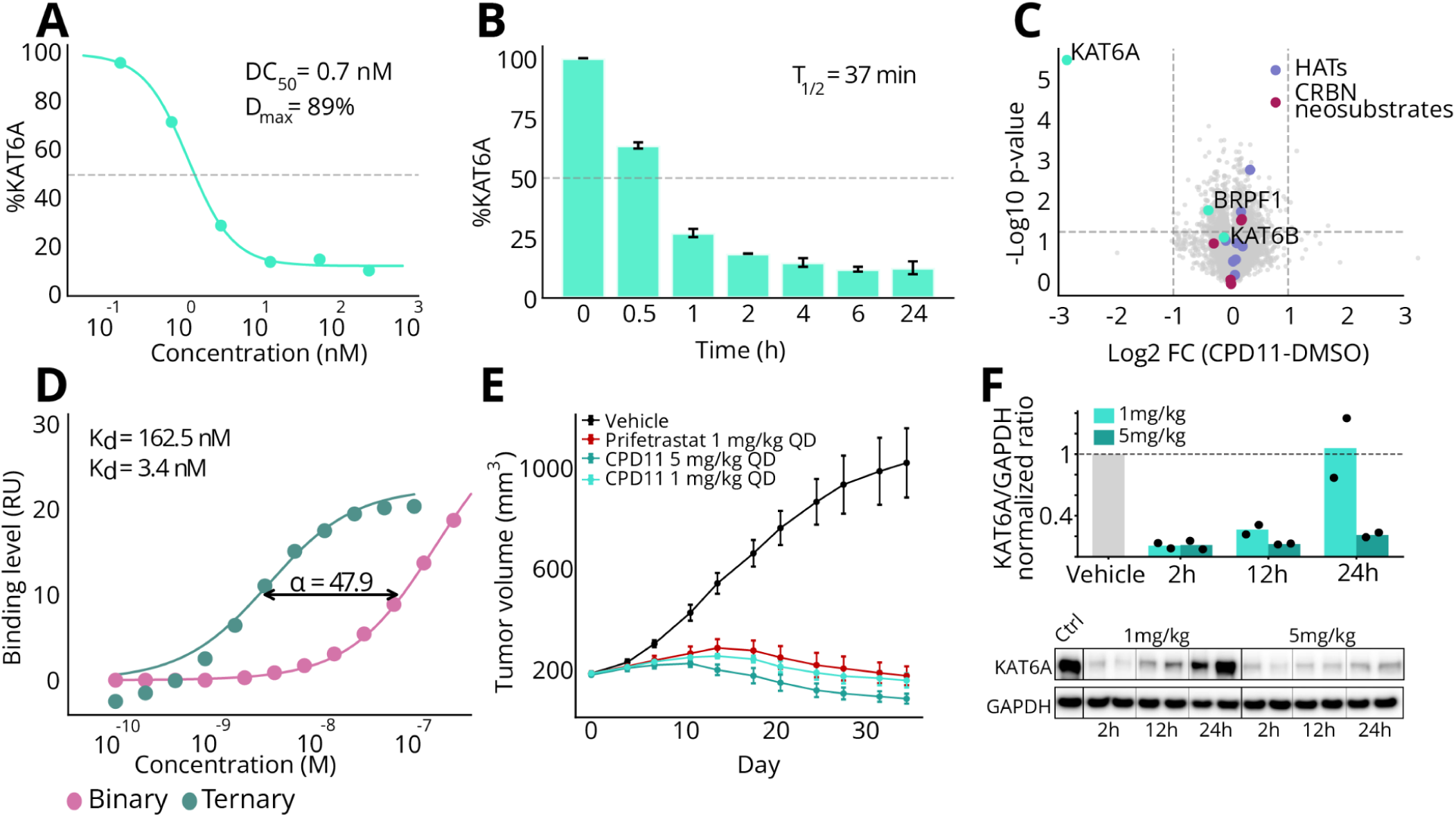
AIMS™ enabled the development of a potent KAT6A degrader. A- Dose-dependent degradation of KAT6A as measured by HiBiT engineered T47D cell line treated with CPD11. B- Time-dependent degradation of KAT6A as measured by HiBiT engineered T47D cell line treated with CPD11 (8nM). C- Global proteomics profile of ZR751 cells treated with KAT6A degrader show a clear degradation profile. KAT6A shows the highest and most significant fold change compared to the entire proteome. No significant changes observed in KAT6B or CRBN neurosubstrates. D- SPR comparing binary and ternary binding affinity of CPD11. E- Efficacy study in ZR-75-1 xenograft mice comparing KAT6A degrader (turquoise shades) and KAT6 inhibitor (red). F- Tumor pharmacodynamics reflects sustained KAT6A degradation. Barplot (top) showing normalized quantification of KAT6A level, measured in duplicates by western blot (bottom).

To evaluate the degradation profile of CPD11, both global and targeted MS proteomics studies were performed. In a global proteomics study, ZR-75-1 breast cancer cells were treated with 8 nM of CPD11 for 6h. KAT6A expression levels were significantly downregulated (Log_2_ Fold change = -2.85, p-value = 3 x 10^-6^). In contrast, the levels of neither KAT6B nor other histone acetyltransferase (HAT) family members were significantly altered, further supporting selectivity towards KAT6A (**Figure 6C**). Additionally, SPR analysis was performed to compare binary and ternary binding affinities. These measurements demonstrated robust ternary complex formation with marked positive cooperativity (ɑ = 47.9, **Figure 6D**).

Motivated by the strong *in vitro* activity of CPD11 and supported by the computational model enabling improved permeability, we evaluated the pharmacokinetics (PK) of CPD11 in orally dosed mice. The compound exhibited good exposure in plasma, maintaining exposure above the equivalent *in vitro* DC50 for ∼16 h, with bioavailability = 27.4%, supporting QD or BID oral administration (**Supplementary Figure S8B**). Together with the *in vitro* data, these results establish CPD11 as a potent degrader with a good pharmacokinetic profile suitable for *in vivo* efficacy testing.

To test the effect of KAT6A degradation *in vivo*, we performed a mouse xenograft efficacy study. Briefly, Balb/c nude mice were implanted with ZR-75-1 ER+ breast cancer cells and orally dosed with CPD11 (1-5 mg/kg daily), compared to vehicle and to the clinical KAT6 inhibitor Prifetrastat (1 mg/kg QD). Remarkably, while the KAT6 inhibitor led to significant tumor growth inhibition, CPD11 (5 mg/kg) showed superior activity, leading to substantial tumor volume reduction (**Figure 6E**). Tumor pharmacodynamic (PD) analysis of mice from CPD11-treated arms revealed sustained reduction of KAT6A levels in the 5 mg/kg arm, supporting sustained drug exposure (**Figure 6F**). In terms of tolerability, CPD11 was generally well-tolerated and no significant changes in the mice body weight were observed (**Supplementary Figure S8C**).

## Discussion

PROTAC development is a multistep and multi-parameter process, and the maximization of one might come at the expense of others^18,49^, highlighting the need for a multi-parameter optimization approach. Here we proposed the first systematic approach to combine structural proteomics, AI models and physics-based methods for rational drug design, and outlined the optimization process of a potent KAT6A degrader using the AIMS™ platform. The incorporation of experimental data into AI models provided biologically-informed structure solving; and its scale and throughput enables effective integration into rapid drug design cycles. One of the main use cases of these capabilities is to model the ternary structure differences between strong and weak degraders, as in the rational design of PROTACs, it is established that even small modifications to a linker can result in large changes in the efficiency of target degradation^50,51^.

Using AIMS™ to study the structures of PRTOAC-dependent ternary complexes provided several structural insights. First, we found that strong KAT6A degraders have a characteristic productive conformation, different from that of weak degraders and not captured by naïve AI models. Second, molecular and metadynamics simulations revealed that active compounds prefer this conformation and are less likely to stabilize an unproductive one; and third, that a PROTAC needs to have a specific exit vector topology in order to enable fast ternary complex assembly. Thus, AIMS-Fold suggested PROTAC-dependent structures that can inform downstream synthesis. Relying on AIMS-Fold insights, AIMS-Rank then utilizes the differentiating structural features for compound ranking. These models ultimately led to the design of a potent KAT6A degrader, which demonstrated significant anti-tumor activity and possessed favorable pharmacological characteristics, making it a promising drug candidate for further development.

During the optimization process, the multi-parameter nature of PROTAC rational design was extensively illustrated. For example, while AIMS-Rank performed well for the majority of compounds, this analysis also highlighted a group of compounds whose degradation potency was not explained by the ternary alone and could be explained by factors such as permeability, solubility, or association and dissociation rates. In another example, we found that favorable Ub geometry is not always associated with better degradation efficiency. Examination of this phenomenon revealed that for some compounds, while they adopt the correct ternary complex geometry upon binding, they suffer from weak/moderate binding affinity. This suggests that a favorable geometry cannot compensate for a compound that fails to reach the target or lacks the baseline affinity to form the complex in the first place.

Although this work has focused on the contribution of our models to optimization of a KAT6A PROTAC, the application of AIMS-Fold to structure solving has been extensively validated in several benchmark cases^43^. However, improving and expanding experimental data could also provide significant improvement for the performance of the models. For example, our structural MS studies are based on recombinant systems. While these studies are highly informative and allow the serial evaluation of multiple compounds, studying complexes in cellular systems would shed additional light on interactions and conformations potentially overlooked by recombinant systems. Other limitations of structural proteomics techniques have been discussed before^52–54^. For example, XL-MS applicability, particularly for PROTAC design, often depends on the prevalence and accessibility of lysine residues on both target protein and E3 ligase, and therefore their absence in the PPI interface might limit its utilization in certain proteins or complexes. On the other hand, overcoming this drawback is partly achieved by the use of HDX-MS derived orthogonal data, and can be further addressed by incorporation of additional crosslinking agents and chemistries that target acidic residues or provide nonspecific mapping^55,56^. A feature common to both XL-MS and HDX-MS techniques is peptide/residue resolution, rather than atomic level resolution provided by other experimental methods. While this does not limit their subsequent integration and contribution to AI models, it might introduce a certain level of noise, as the protection patterns or distance between some residues might be affected by residues in their proximity. Mitigating this potential noise can be done by external validation using complementary approaches, and has been extensively shown previously both for HDX-MS^57,58^ and XL-MS^59,60^.

In this case study, our tools were successfully used to develop a model based on productive conformations; and while other models were also developed in order to address the various layers of PROTAC development and showed overall good performance in the case of KAT6A, future work will address additional parameters for optimization and determine the unique predictive contribution of these models to additional targets. Nevertheless, the concepts and tools demonstrated here are not unique to KAT6A per se. While one cannot guarantee in advance what factors are most likely to contribute in a future drug discovery program, these approaches can be readily applied to additional targets, as well as additional proximity inducing modalities such as molecular glue degraders (MGDs), RIPTACs, and others^61^. Beyond the application to different modalities, the platform is also expected to benefit from integration of native MS data, enrichment of XL-MS data as mentioned above, and further optimization of the translation of structural MS data into the computational models.

The contribution of AI to drug discovery remains a subject of substantial discussion, as well as its impact on translational and therapeutic success. In a recently published review, Coley and colleagues extensively outline the challenges of AI in drug discovery^62^. They suggest a granular breakdown of the challenges into different aspects (Chemistry, Pharmacology, Structure, and Energy); and claim that the translational benefit of AI would eventually manifest in solving specific, well-defined problems. Consistent with this approach, the various modules of our platform address well-defined PROTAC optimization aspects. Our experimentally grounded models deliver higher biological fidelity compared to naïve AI alongside higher speed and scalability compared to conventional structural techniques. Collectively, they enable a systematic approach to a multi-faceted challenge, bringing the promise of AI closer to reality and paving the way for rapid discovery of best-in-class drugs.

## Methods

### Hydrogen-deuterium exchange mass spectrometry (HDX-MS)

MYST domain of human KAT6A (501-784) was purchased from Crelux (WuXi AppTec company, China). The construct was expressed and purified from insect cells and validated by SEC. Full length CRBN (1-442)/ DDB1 (1-1140)/ DDA1 (1-102), expressed and purified from Baculovirus, was purchased from Viva biotechnology (China). PROTACs were synthesised byWuXi AppTec company (China)

Briefly, the two proteins and PROTAC compounds were incubated in the ratio of CRBN : PROTAC : KAT6A (1 : 1.5 : 1) for ternary complex analysis. For CRBN : PROTAC and KAT6A : PROTAC binary complex analysis, the samples were incubated in the ratio of 1:1.5. APO state KAT6A and CRBN served as controls. HDX of KAT6A and CRBN protein and their complexes was initiated by 5-fold dilution in a deuterated buffer containing 50mM HEPES (pH/pD 7.6), 100mM NaCl and 2mM MgCl2. The deuteration reactions were carried out for set timepoints of 30 sec, 15 min and 2hrs at 20 °C. All data points were measured in triplicate and the labelling reaction was quenched by adding a quenching buffer containing 1M Glycine, 4M Urea, 250mM TCEP, pH 2.3. All samples were handled using the automated platform (AffiPro HDX workstation). The protein samples were digested online using co-immobilized Nepenthesin-2/pepsin column (AffiPro) at a flow rate of 0.2mL/min. Peptides were separated using a UPLC system consisting of an Agilent 1290 Infinity II pump and 1260 isocratic pump (Agilent Technologies, Santa Clara, CA, USA). A Phenomenex Luna Omega Polar C18 analytical column (100 × 1 mm) was used for peptide separation and mass spectra were obtained with timsToF Pro (Bruker Daltonics). To minimize the loss of deuterium, the LC system was refrigerated to 0 °C.

The mass spectrometer operated in MS mode with a 1 Hz data acquisition rate. For peptide identification, the same LC-MS system was used, but the mass spectrometer operated in data-dependent MS/MS mode using PASEF. The LC-MS/MS data were searched using MASCOT (v. 2.7, Matrix Science) against a customized database combining sequences of KAT6A and CRBN. Search parameters were set as follows: no-enzyme, no modifications allowed, precursor tolerance 10 ppm, fragment ion tolerance 0.05 Da, decoy search enabled, FDR < 1%, IonScore > 20, and peptide length > 5. Acquired LC-MS data were peak-picked and exported in DataAnalysis (v. 5.3, Bruker Daltonics) and further processed by the DeutEx software (Bruker Daltonics). Average deuteration per amino acid was calculated based on the workflow described by Trcka et al.^62,63^. Percent deuteration values are reported as mean ± standard deviation (SD) from three technical replicates.

### Downstream analysis

The estimation of individual amino acid uptake was performed as described in Trcka *et al*^63^ with an addition of a standard deviation based peptide filtering. Briefly, individual peptides were removed if their uptake SD was higher than 2.5 standard deviations compared to the mean peptide SD of the sample. Then, the uptake of a given amino acid was calculated as the weighted mean of all overlapping peptides covering that residue. The weight applied for each peptide was inversely proportional to its length to ensure higher contribution of shorter peptides, higher resolution peptides.

To quantify the predicted interaction interfaces, the solvent-accessible surface area (SASA) of each ternary model was calculated. Then, the ΔSASA of each residue upon complex formation was calculated by subtracting the SASA of the full ternary complex from its SASA in the isolated protein chain.

To assign an HDX based score for each predicted ternary complex, KAT6A and CRBN residues were categorized into hotspots (Ternary-Apo protection > 4%) and coldspots (Ternary-Apo protection < 4%). Each ternary model was rewarded the burial of experimentally buried regions by summing the product of the HDX based protection and the ΔSASA for all hotspot defined residues. Burial of cold spots was used to penalize false positive interfaces. Specifically, the penalty subtracted for each buried cold spot was calculated as the product of its ΔSASA, the average absolute HDX magnitude of all hotspots and a class balancing weight (the ratio of total number of hotspot residues to coldspot residues).

### Cross linking mass spectrometry (XL-MS)

MYST domain of human KAT6A (501-784) and mini CRBN (CRBN(41-187)-GSG-CRBN (249-426), expressed and purified from E. coli, was purchased from Viva biotechnology (China). Recombinant KAT6A protein and PROTAC were incubated in the ratio of 1 : 1.5 for 30 minutes at room temperature in the reaction buffer (50 mM HEPES, 100 mM NaCl, pH 7.6, 2mM MgCl2). Following this, a molar equivalent of CRBN was added to the samples and further incubated for 15 minutes at room temperature. PROTAC compounds were replaced with DMSO to serve as a control. Samples were crosslinked with an equimolar mixture of Bis(sulfosuccinimidyl) suberate (BS3) and Disuccinimidyl suberate (DSS) for 30 minutes (0.5mM final conc). The reaction was quenched by incubating the sample with a 1M ammonium bicarbonate buffer (100mM final concentration) for 15 minutes at room temperature.

Following the crosslinking procedure, protein samples were mixed with 2 volume equivalents of denaturing buffer containing 0.2% n-dodecyl-β-D-maltoside (DDM), 10mM tris(2-carboxyethyl)phosphine (TCEP), 40mM chloroacetamide (CAA) and 200mM triethylammonium bicarbonate (TEAB) buffer pH 8.5 and incubated at 55°C for 30 min. Proteolytic digestion was initiated by addition of 1 volume equivalent of 50 mM ABC (pH 8.0) containing trypsin (enzyme:protein, 1:12.5 w/w) and Lys-C (enzyme:protein , 1:25 w/w), followed by overnight incubation at 37°C. Resulting peptides were desalted with Oasis HLB cartridges (Waters Corporation, Milford, MA, USA) prior to LC–MS/MS analysis.

Peptide samples were analyzed using a Vanquish Neo nanoLC system coupled to an Orbitrap Exploris 480 mass spectrometer operating in positive ion mode. Analytical separation was carried out on an Aurora Ultimate C18 analytical column (25 cm × 75 µm ID, AUR4). Data was acquired on the Orbitrap Exploris 480 in data-dependent acquisition (DDA) mode with a Top7 method. Full MS survey scans were acquired in the Orbitrap over the m/z 380–1200 range at a resolution of 60,000. MS/MS spectra were acquired for precursors with charge states 4–8, using a 2 m/z isolation window and HCD fragmentation with stepped normalized collision energies of 25, 30, and 35. MS/MS scans were collected at 30,000 resolution.

Crosslinked peptide identification was performed for each sample using MeroX (version 2.0.1.4). The crosslinker composition was set to C₈H₁₀O₂ and monoisotopic mass of 138.06808 Da. Digestion enzyme was set as trypsin, allowing up to three missed cleavages, and peptides with lengths between four and thirty amino acids were retained. Carbamidomethylation of cysteine was set as a static modification, and oxidation of methionine was included as a variable modification. Crosslinking specificity was restricted to lysine–lysine (K–K) pairs. Mass tolerances were set to ±5 ppm for precursor ions and ±10 ppm for fragment ions. Data were analyzed in Quadratic Mode, using a minimum peptide score of 1 and a score cutoff of 20. Only crosslinks passing a 1% false discovery rate (FDR) were retained for further analysis, and a signal-to-noise ratio threshold of 1.0 was applied to improve confidence in peptide identification.

For crosslink unification across samples, the raw MS files were converted to MZML using ProteoWizard MSConvert and the results of all MeroX output files were aggregated. Redundant spectral matches (±0.01 m/z, RT ±60 seconds) were clustered. Crosslink clusters that were detected in fewer than 3 samples or with a median MeroX score <60 were filtered out. The median RT and m/z of the remaining clusters were calculated and were used to extract the intensity from each MZML file (dynamic m/z tolerance of 10 ppm and RT ±8 seconds). Finally, the intensities of the same crosslinked peptides originating from different charges and RT were summed.

To account for technical differences between samples, a total intensity normalization was performed and the data were log_2_ transformed(log_2_ ((intensity / total_intensity) * constant + 1)). The average of all technical repeats was calculated for each crosslinks under each PROTAC treatment. For detecting differential inter-protein crosslinks, the log_2_ fold change between the average of strong (<50 nM) and weak compounds (>1000nM) was calculated for each crosslink.

To assign an XL-MS based score for each predicted ternary complex, the distance between the lysine alpha-carbons of each crosslink of interest was calculated. The distance was converted to a satisfaction probability using a sigmoidal decay function where distances <15 Å received a probability of 1 and >15 Å were penalized via a sigmoid curve with an inflection point at 25 Å. Next, a weight was assigned to each crosslink by the product of its overall intensity (average across all samples) and the log_2_ fold change between potent and inactive compounds. The product of the satisfaction score and the weight was calculated for each crosslink. The sum of these products for all differential crosslinks defines the ternary complex score. High positive values correspond to structures representing potent compounds and low negative scores to ones representing inactive ones.

### Ubiquitinated peptide enrichment and MS analysis

6×0^6^ ZR751 cells were plated in 10cm plates with RPMI-1640 growth medium supplemented with 10% fetal bovine serum (FBS), 1 mM sodium pyruvate, 2 mM L-glutamine, and 1% penicillin–streptomycin. Cells were treated for 30 minutes with 8nM compound along with 10uM MG-132 proteasome inhibitor. Samples were harvested for MS analysis after 3 washes with PBS.

Cells were lysed in buffer containing 0.2% n-dodecyl-β-D-maltoside (DDM), 10 mM tris(2-carboxyethyl)phosphine (TCEP), 40 mM chloroacetamide (CAA), and 200 mM triethylammonium bicarbonate (TEAB), pH 7.8. Lysates were heated at 95 °C for 10 min and sonicated. Following this, samples were incubated for 30 minutes at room temperature with Benzonase in the presence of 2mM MgCl₂ to rescue sample viscosity. The resulting protein samples were subjected to a two-stage digestion protocol. First, trypsin was added to the samples and incubated for 2–3 h at 37 °C. Samples were then centrifuged at 17,000 × g, and the supernatant was transferred to a fresh tube. The remaining pellet was resuspended in 9 M urea, followed by overnight digestion with trypsin at room temperature. Digestions were performed using Lys-C (1:100, w/w) and trypsin (1:50, w/w) and quenched by addition of trifluoroacetic acid (TFA) to a final concentration of ∼1%. The peptides from supernatant and pellet fractions were subsequently combined and desalted with C18 cartridges.

Immunoaffinity enrichment of acetylated peptides was performed with PTMScan HS K-epsilon-GG Remnant Magnetic Immunoaffinity Beads (Cell Signalling Technologies #34608) according to manufacturers instructions with modifications. Briefly, about 1mg of vacuum dried peptides were resuspended in pH adjusted 1× HS IAP Bind Buffer #1. Antibody-conjugated beads were washed four times with ice-cold PBS and incubated with peptide samples for 2 h at 4 °C with gentle agitation. Following incubation, beads were separated magnetically and washed five times with HS IAP Wash Buffer and twice with LC–MS-grade water. Bound peptides were eluted with 0.15% TFA, desalted with Oasis HLB cartridges (Waters Corporation, Milford, MA, USA) and resuspend with 5ul of buffer containing 3% ACN, 97% H2O, 0.1% TFA, 0.015% DDM.

Peptides were separated over a chromatography 194.5 min gradient and analyzed using an Orbitrap Exploris 480 mass spectrometer (Thermo Fisher Scientific). Data were acquired using a data-independent acquisition (DIA) strategy where full MS scans were acquired at a resolution of 120,000 (at m/z 200), over a mass range of m/z 375–1500 and MS/MS spectra were acquired at a resolution of 30,000. Fragment ions were generated using higher-energy collisional dissociation (HCD) with stepped normalized collision energies of 25%, 27.5%, and 30%.

Raw mass spectrometry data files were processed using DIA-NN software (version 1.8.1). To identify ubiquitinated peptides, we used the library-free search option, where an in silico spectral library was generated from a human FASTA database (UniProt 2024-03). The search allowed for trypsin peptides with a maximum of two missed cleavages. For fixed modification we used carbamidomethylation of cysteine, and for variable modifications oxidation of methionine, protein N-terminal acetylation, and ubiquitin on lysine residues were used. Match Between Runs (MBR) feature was enabled. Peptide precursor and protein-level false discovery rates (FDR) were strictly controlled at 1% using DIA-NN’s target-decoy strategy.

Quantitative ubiquitination data was processed using the log2-transformed intensity values for each detected site. Missing values were imputed using values derived from the lower tail of each individual sample’s distribution. Since the PROTAC directly engages the KAT6A MYST domain, we focused our analysis exclusively on all ubiquitinated lysines within this targeted region. To quantify the overall ubiquitination shift shown in Figure 2A, we calculated the sum of the log2 intensity differences between the potent PROTAC-treated samples and the DMSO controls across all detected MYST domain sites.

### Structural modeling with structural MS constraints (AIMS-Fold)

For ternary complex structure prediction, we used AIMS-Fold^43^, a diffusion-based generative model built on Boltz-2. It integrates structural proteomics data into the generation process using inference-time guidance, overcoming the limitations of traditional post-hoc filtering. Specifically, the model translates mass spectrometry data into two types of active constraints. Cross-linking mass spectrometry (XL-MS) data is converted into spatial constraints, using both positive constraints for proximate residues and negative constraints for residues far apart. Additionally, hydrogen-deuterium exchange (HDX-MS) data is incorporated using a differentiable burial potential, capturing proximate interactions and interface flexibility.

### Meta dynamics for stabilization ranking

Protein structures were obtained using structure prediction tools with distance constraints to ensure consistent starting positions for different compounds. The structures were prepared using the Schrodinger Maestro 13.7 with protonation states set to pH 7.4.

The systems were solvated and neutralized using theMolcube platform. Each system was placed in a truncated octahedral box with 1.5nm margin, solvated with TIP3P water model and neutralized by adding 0.15M NaCl. The system parametrization was performed with CHARMM36 force field for the proteins and OpenFF force field for the ligands.

Each system underwent energy minimization and NVT equilibration to stabilize the system to 300K using Gromacs 2021.7. Then, GPU accelerated Well-Tempered Metadynamics was performed for each of the systems using Gromacs patched with Plumed. Three collective variables (CVs) were chosen to bias the sampling towards exploring conformational space that was acquired using multiple structure prediction tools with different constraints.

The conformational space was constructed from two types of movements -hinge movement and rotational swivel. To explore the hinge movement, we defined two CVs -one angle CV and one distance CV. The rotational movement was monitored using a dihedral CV, defined as the angle between two planes passing through the centers of each protein. Each system was run in six independent replicates with each run lasting up to 150ns. Walls were defined to break the simulation upon ligand disengagement from either one of the proteins. Gaussians were deposited every 1ps with an initial height of 2 KJ/mol. The gaussian widths (sigma) were set in the range of 0.1-0.15 for all three CVs with a bias factor of 15.

The resulting Free Energy Surfaces (FES) were analyzed and projected for each two CVs combination. Dividing the projections of CV1 and CV2 into quadrants allowed to prioritize the compounds based on their residency in Q4 (bottom right).

### Predicted activity based on the active conformation (AIMS-Rank)

Metadynamics simulations indicated that potent compounds preferentially stabilize in the upper left quadrant (Q1) of the free energy landscape (Figure 2A), corresponding to the AIMS-Fold-predicted active conformation. Positing that higher binding affinity within this active state correlates with improved degradation, we developed a targeted scoring approach. Using AIMS-Fold guided by active-state cross-linking mass spectrometry (XL-MS) constraints, we generated an ensemble of 20 distinct ternary conformations for each compound. The overall activity score was subsequently defined as the median predicted ternary affinity of the conformations falling within the Q1 basin, with individual model affinities calculated using a modified version of Boltz-2 adapted for single-model scoring.

### Computational Prediction of Target Ubiquitination

To capture the conformational flexibility of the ligase machinery, we constructed an ensemble of 9 full E3-E2 ligase models, incorporating both ring-forming and open DDB1 states following the methodology described by Bai et al. The assembled ligase complexes (comprising CRBN [PDB ID: 4TZ4], DDB1 [PDB IDs: 2B5L, 3E0C, 3EI3, 3EI4, 3I8E, 4A0L, 4E54, 4TZ4, 6PAI], CUL4A [PDB ID: 2HYE], Rbx1 [PDB ID: 6TTU], NEDD8 [PDB ID: 6TTU], E2 [PDB ID: 5FER], and ubiquitin [PDB ID: 6TTU]) were subsequently energy-minimized using the Schrödinger software suite 2024-02. This was performed using the OPLS force field with a standard restrained minimization protocol to relieve local steric clashes and optimize sidechain geometries.

For each predicted ternary conformation (CRBN-PROTAC-POI) in our generated ensemble, we performed a rigid-body superimposition onto the ligase ensemble. The CRBN chain of the ternary complex was aligned to the CRBN chain of each of the 9 minimized ligase models by aligning their alpha-carbon (Cα) backbone atoms. This rotational and translational matrix was then applied to the target protein, positioning it within the ligase active site. To filter out physically impossible geometries, we evaluated the newly assembled full complexes for severe steric clashes. We calculated the pairwise distances between the heavy atoms of the target protein and the heavy atoms of the ligase engine components (CUL4A, Rbx1, NEDD8, E2, and DDB1). Models were flagged as clashing if more than 20 target heavy atoms fell within a 2.0 Å radius of the ligase machinery. Ternary conformations that exhibited clashes across all 9 ligase models were classified as unproductive and discarded from further analysis.

To quantify the ubiquitination potential of viable ternary conformations, we focused on three target lysines. These residues were selected based on our quantitative proteomics assay, as they exhibited the strongest ubiquitination signals with a >6 log2 fold change when comparing potent compound treatment to DMSO controls. For every viable, non-clashing model, we calculated the Euclidean distance from the C-terminal carbonyl carbon of ubiquitin (Gly76) to the sidechain epsilon-nitrogen (Nζ) of the three target lysines. To assign a single proximity metric to each ternary conformation, we first identified the minimum spatial distance achieved for each target lysine across its viable ligase models. We then calculated the average of these minimum values across all three target lysines, yielding a final metric that represents how favorably the specific ternary conformation positions the target for ubiquitin transfer.

### Unbinding kinetics and exit topology prediction

To investigate the unbinding kinetics and exit topology of the PROTAC-target complexes, we employed well-tempered metadynamics (WT-MetaD). Critical intermolecular interactions between the target protein and the PROTAC were selected as collective variables (CVs) for the biasing potential. This biasing strategy encouraged the PROTAC to explore and identify the lowest energy exit pathway from the binding pocket. A COMMITTOR function was defined to automatically terminate the simulation once the PROTAC had fully dissociated from the target. This method measures the energetic barrier associated with ligand unbinding. To ensure statistical robustness, ten independent WT-MetaD simulation replicates were performed for each evaluated PROTAC.

Following the unbinding simulations, the generated trajectories were post-processed and reweighted using the PLUMED 2.9.1 plugin. The systems were reweighted along three additional structural CVs: the linker exit vector angle, the linker buried surface area (BSA), and the dihedral torsion angle at the linker attachment point. Free energy surfaces (FES) were subsequently reconstructed based on these reweighted CVs. Analysis of these energy landscapes revealed a topological difference between strong and weak degraders, which was characterized by the orientation of the linker exit vector and the extent to which the linker was buried against the protein surface.

### Conformational sampling simulations

To evaluate the membrane permeability and chameleonic behavior of the PROTACs, molecular dynamics (MD) simulations were conducted in both polar (water) and apolar (chloroform) environments to mimic the aqueous cellular solvent and the lipid bilayer, respectively, as shown previously^48,63^. Initial 3D structures were generated using the Maestro suite (Schrödinger) and prepared via LigPrep before being exported in SDF format. Each system was solvated in a truncated octahedron box, maintaining a minimum buffer distance of 10 Å between the PROTAC and the periodic boundaries. For the apolar environment, chloroform was added to achieve the experimental density of 7.4 molecules/nm³. Prior to the production runs, the systems underwent an equilibration protocol consisting of energy minimization to resolve steric clashes, followed by canonical (NVT) and isothermal-isobaric (NPT) ensemble simulations to equilibrate the system to the target temperature and pressure, respectively. Production MD simulations were then performed in triplicate for 100 ns per system in each solvent. Following the simulations, the resulting trajectories were centered to correct for periodic boundary crossings and ensure the PROTAC remained in the middle of the box. To quantify chameleonicity, specifically the ability of the PROTAC to adopt an extended state in water and a compact, polarity-shielded conformation in the membrane, two key structural metrics were calculated using the GROMACS [2021.7] software package: the radius of gyration (Rg) to assess molecular size and compaction (via gmx gyrate), and the solvent-accessible 3D polar surface area (3D PSA), defined as the SASA of nitrogen, oxygen, and sulfur atoms (via gmx sasa) to evaluate the exposure of polar groups^64^.

### HiBiT-based KAT6A degradation

Compound-induced degradation of KAT6A was assessed by quantifying cellular KAT6A protein levels using a HiBiT reporter system. Experiments were performed in T47D cells engineered to express HiBiT-tagged KAT6A. Cells were cultured at 37 °C in a humidified atmosphere containing 5% CO₂, using RPMI-1640 medium supplemented with 10% FBS, 0.008mg/ml insulin and 1% penicillin–streptomycin. For the assay, cells were plated in 96-well white microplates at 20,000 cells per well. Test compounds, prepared in DMSO, were applied as a six-point serial dilution. Following a 6-hour incubation, intracellular KAT6A levels were determined using the Nano-Glo HiBiT Lytic Detection reagent according to the supplier’s protocol. Luminescence was recorded on a multimode plate reader (BMG CLARIOstar). Raw signal intensities were background-subtracted and normalized relative to DMSO-treated controls. DC50 was calculated by fitting the data to a four-parameter sigmoidal dose-response (variable slope) using GraphPad Prism.

### *In vitro* viability assay

ZR-75-1 Human cancer cell lines were maintained under standard cell culture conditions at 37 °C in a humidified atmosphere containing 5% CO₂. Cells were cultured in RPMI-1640 medium supplemented with 10% FBS and 1% penicillin–streptomycin. Test compounds were dissolved in DMSO to prepare stock solutions. Working solutions were prepared by dilution in assay medium immediately before treatment. The final DMSO concentration in all assay wells was maintained at 0.03%. Cells were counted using Countess 3 Thermo Fisher Scientific, and adjusted to the desired seeding density (2000 cells/well). Cells were seeded in 100 µL assay medium. Blank wells contained medium only. Test compounds were dispensed into assay plates in 10 points, 4 fold serial dilution, starting at 3000nM down to 0.01nM. After compound addition, plates were returned to the incubator and maintained for either 10 days. Culture medium was replaced on day 5 and fresh compound was re-added at each medium change. Cell viability was quantified using a luminescence-based ATP detection assay according to the manufacturer’s instructions. Briefly, assay plates and reagents were equilibrated to room temperature. 60ul of CellTiter-Glo reagent was added directly to each well to induce cell lysis. Plates were mixed on a shaker for approximately 2 minutes and incubated at room temperature for an additional 10 minutes to allow luminescent signal stabilization. Luminescence was measured using a multimode plate reader. Raw luminescence values were used as a measure of relative cell viability and were normalized to vehicle-treated controls. IC50 was calculated by fitting the data to a four-parameter sigmoidal dose-response (variable slope) with top constraint to 100.

### Surface plasmon resonance (SPR)

SPR measurements were performed on a BiacoreTM 8K instrument (Cytiva Life Sciences). For all experiments, binding data were analyzed using Biacore Insight Evaluation Software. Sensorgrams were double-referenced by subtracting the response from reference flow cell 1 and a buffer blank, and solvent correction was applied using an 8-point DMSO standard curve. Binding curves were fitted with a 1:1 kinetic or affinity model to derive the association rate constant (ka), dissociation rate constant (kd), and equilibrium dissociation constant (KD).

For binary KAT6A binding, the sample compartment and flow cell were maintained at 20°C. A Sensor Chip NTA was equilibrated in running buffer containing 10 mM HEPES (pH 7.5), 150 mM NaCl, 0.25 mM TCEP, and 0.005% Tween-20. The chip surface was preloaded with 350 mM EDTA for 60 s, followed by 0.5 M NiCl2 for 60 s, and then activated with a 50% EDC/50% NHS mixture for 420 s. N-terminal His-tagged KAT6A was immobilized on flow cell 2 at 3 µg/mL in running buffer at 10 µL/min to a level of 1500-1700 response units (RU). The running buffer was supplemented with 2% DMSO, and compounds were prepared as 2-fold serial dilutions across 9 concentrations. After 30 startup cycles, compounds were injected using single-cycle kinetics with a contact time of 90 s per concentration and a dissociation time of 1200 s at a flow rate of 30 µL/min.

For binary CRBN/DDB1 binding, the sample compartment and flow cell were maintained at 20°C. A streptavidin (SA) sensor chip was equilibrated in running buffer containing 10 mM HEPES (pH 7.0), 150 mM NaCl, 0.5 mM TCEP, and 0.005% Tween-20. The chip surface was conditioned by three injections of 1 M NaCl and 50 mM NaOH for 60 s each. Avi-tagged CRBN/DDB1 was captured on flow cell 2 at 60 µg/mL for 1200 s at 5 µL/min to an immobilization level of 5700-6000 RU. The running buffer was supplemented with 2% DMSO, and compounds were prepared as 2-fold serial dilutions across 11 concentrations. After 20 startup cycles, compounds were injected using multi-cycle kinetics with a contact time of 90 s and a dissociation time of 600 s at a flow rate of 30 µL/min.

For ternary complex SPR measurements of CRBN/DDB1, KAT6A, and small molecules, the flow cell was maintained at 20°C and the sample compartment at 10°C. A streptavidin (SA) sensor chip was equilibrated in running buffer containing 10 mM HEPES (pH 7.0), 150 mM NaCl, 0.5 mM TCEP, and 0.005% Tween-20. The chip surface was conditioned by three injections of 1 M NaCl and 50 mM NaOH for 60 s each. Avi-tagged CRBN/DDB1 was captured on flow cell 2 at 1 µg/mL and 5 µL/min to an immobilization level of 200 RU. The running buffer was supplemented with 2% DMSO, and compounds were prepared as 2-fold serial dilutions across 11 concentrations. After 20 startup cycles, compounds were injected in the presence of a constant 100 nM KAT6A using multi-cycle kinetics with a contact time of 90 s and a dissociation time of 600 s at a flow rate of 30 µL/min.

### Pharmacokinetics

Plasma pharmacokinetics of the compound were evaluated in male CD-1 mice (6-10 weeks old; n = 3 per group). Animals were acclimated for at least 3 days before study initiation and were maintained under standard environmental conditions with ad libitum access to food and water, except where fasting was required. For the oral dosing group, animals were fasted overnight prior to dosing and until 4 h post-dose. All procedures were conducted in accordance with institutional animal care and use guidelines at WuXi AppTec and under non-GLP conditions. The compound was administered either as a single intravenous bolus dose at 1 mg/kg or by oral gavage at 5 mg/kg. The vehicle for both routes consisted of 5% DMSO, 10% Solutol, and 85% water. All formulations were prepared within 4 h prior to dosing, and aliquots were collected for dose concentration verification by LC/UV or LC-MS/MS. For the intravenous group, animals received a dose volume of 5 mL/kg at a target formulation concentration of 0.2 mg/mL. For the oral group, animals received a dose volume of 10 mL/kg at a target formulation concentration of 0.5 mg/mL. Dose volumes were calculated based on body weight measured on the day of dosing. Intravenous administration was performed via tail vein bolus injection or indwelling cannula, and oral dosing was performed by gavage. Blood samples were collected into K2-EDTA tubes via the saphenous vein or another suitable blood collection site. In the intravenous group, plasma samples were collected at 0.083, 0.25, 0.5, 1, 2, 4, 8, and 24 h post-dose. In the oral group, plasma samples were collected at 0.25, 0.5, 1, 2, 4, 8, and 24 h post-dose. Each blood sample was approximately 0.025 mL and was kept on wet ice until processing. Samples were centrifuged at 3200 × g for 10 min at 2-8°C within 1 h of collection, and plasma was transferred into labeled polypropylene tubes and stored at -60°C or below until bioanalysis. Animals were monitored clinically on the dosing day and at sample collection time points, and all animals were euthanized by CO2 inhalation at the final pharmacokinetic time point. Plasma concentrations of the compound were quantified using a non-GLP LC-MS/MS method developed for analysis in biological matrices. Calibration curves included at least 6 non-zero standards including the lower limit of quantification (LLOQ). For runs containing more than 12 samples, quality control samples at low, medium, and high concentrations were included. Bioanalytical acceptance criteria required that at least 75% of calibration standards fall within ±20% of nominal values for biofluid samples, and at least 67% of quality control samples fall within ±20% of nominal values, with at least half of the quality controls passing at each concentration. The target LLOQ was ≤3 ng/mL. Plasma concentration-time data were analyzed by noncompartmental analysis using Phoenix WinNonlin version 6.3 or higher. Pharmacokinetic parameters included, as applicable, clearance (CL), volume of distribution at steady state (Vdss), and C0 for intravenous administration; Cmax, Tmax, and oral bioavailability (%F) for extravascular administration; and terminal half-life (T1/2), AUC0-t, AUC0-inf, MRT0-t, and MRT0-inf for all dosing routes

### In vivo efficacy

ZR-75-1 human breast cancer cells were maintained in RPMI-1640 medium supplemented with 10% fetal bovine serum and 1% penicillin-streptomycin at 37 °C in a humidified atmosphere containing 5% CO₂. Cells were passaged twice weekly and harvested during the exponential growth phase for tumor inoculation. Female BALB/c nude mice (6-8 weeks old, 18-22 g) were obtained from a commercial vendor. Animals were acclimated for approximately one week prior to study initiation and housed under specific pathogen-free conditions with a 12 h light/dark cycle, temperature of 20-26 °C, and humidity of 40-70%. Food and water were provided ad libitum. To support estrogen-dependent tumor growth, a sustained-release β-estradiol pellet (0.36 mg, 90-day release) was implanted subcutaneously 2-3 days prior to tumor inoculation. Mice were inoculated subcutaneously in the right flank with 1 × 10⁷ ZR-75-1 cells suspended in 0.2 mL PBS:Matrigel (1:1). Tumor growth was monitored until mean tumor volume reached approximately 200 mm³, at which point animals were randomized into treatment groups using a block design based on tumor volume. Mice (n = 4-5 per group) were treated with vehicle, a reference inhibitor, or test compounds via oral (p.o.) or intraperitoneal (i.p.) administration. Treatments were administered once daily (QD) or twice daily (BID; 12-hour interval) for up to 42 days depending on group. Treatment arms included vehicle control, reference inhibitor control, and multiple test compound arms administered at different dose levels, schedules, and routes. These included oral and intraperitoneal dosing regimens ranging from 1 to 30 mg/kg, depending on the compound and study arm. Selected groups were terminated at predefined time points for pharmacokinetic (PK) and pharmacodynamic (PD) analyses. A subset of animals demonstrating tumor regression was monitored during a dose-free observation period until study end. Tumor volumes were measured twice weekly. Animals were monitored daily for clinical signs and weighed three times per week. Humane endpoints included body weight loss of at least 20%, tumor volume of at least 2000 mm³, or signs of morbidity including lethargy, impaired mobility, or ulceration. Animals meeting these criteria were euthanized by CO₂ inhalation.

## Author Contributions

N.S., K.P., and E.S. designed the study and supervised the work. Computational structural modeling and data analysis were performed by A.Sht., N.S., S.B., and M.R.S. Experimental studies were conducted by Y.K., I.A., G.O., and N.C. Proteomics studies were conducted by A.She. and D.K. The manuscript was written by G.A., N.S., and S.B., and reviewed by K.L., Y.B.S.G., and all other authors.

## Competing interests

All authors are employees and shareholders of Protai Bio, Ramat Gan, Israel.

## Acknowledgements

The authors wish to thank Ian Churcher for fruitful discussions and extensive and critical review of the manuscript.

