## Supplementary Information for "Leveraging AI and structural proteomics for rational design of a KAT6A degrader"

**Supplementary Figure S1:** XL-MS data shows PROTAC-dependent crosslinks (linked to Figure 1A)

**Supplementary Figure S2:** Comparison of deuterium uptake between ternary and binary complexes (linked to Figure 1B)

**Supplementary Figure S3:** Metadynamics simulations of additional KAT6A degraders (linked to Figure 2B)

**Supplementary Figure S4:** Association rate accounts for AIMS-Rank prediction inaccuracy

**Supplementary Figure S5:** Metadynamics simulations show exit vector angle differentiates between compounds (linked to Figure 4B)

**Supplementary Figure S6:** Chameleonic behavior explains differences in compound bioavailability (linked to Figure 5)

**Supplementary Figure S7:** Multi-parameter optimization highlights potential degrader leads

**Supplementary Figure S8**: CPD11 evaluation *in vitro* and *in vivo* (linked to Figure 6)

**Supplementary Video 1:**
Molecular dynamics simulation shows the transitioning between unproductive and productive states. In the beginning of the simulation CRBN (gray) covers Region B (left) of KAT6A (brown), representing the unproductive conformation. As the simulation progresses, CRBN shifts its position to protect Region A (right), adopting the productive conformation. The PROTAC is colored in yellow, and all trajectory frames are aligned to KAT6A. Region A and B shift between dark and lighter blue, mimicking the HDX-MS patterns of inactive and potent compounds.

**Supplementary Table 1:**
List of 184 generated conformations and whether they introduced steric clashes with each of the nine E2-Ub models.

---

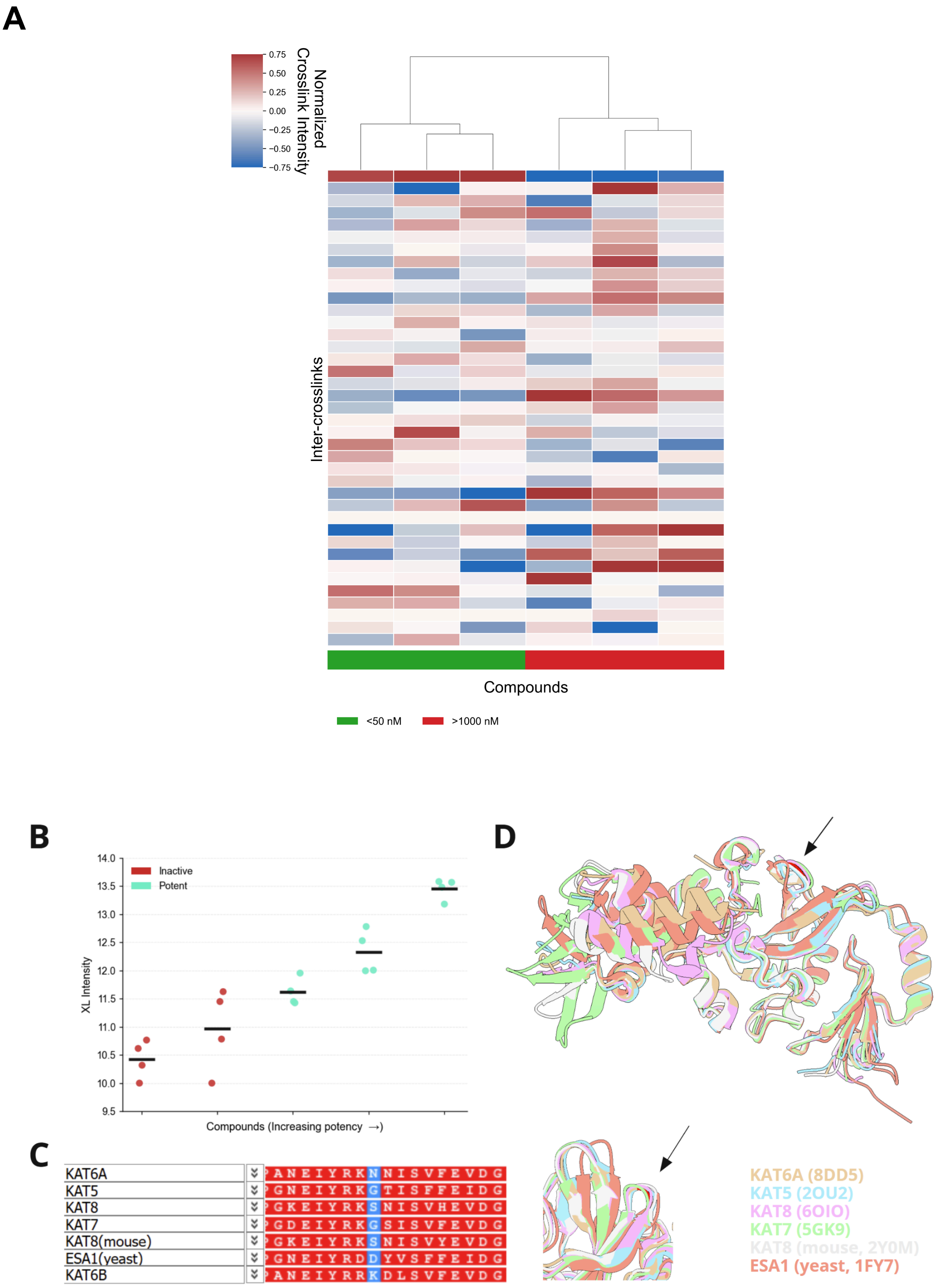

A
0.75
0.50
0.25
0.00
−0.25
−0.50
−0.75
Normalized Crosslink Intensity
Inter-crosslinks
Compounds
<50 nM
>1000 nM
B
14.0
13.5
13.0
12.5
12.0
11.5
11.0
10.5
10.0
9.5
Inactive
Potent
XL Intensity
Compounds (Increasing potency →)
C
KAT6A PANEIYRKNNISVFEVDG
KAT5 GNEIYRKGTISFFEIDG
KAT8 GKEIYRKSNISVHEVDG
KAT7 GDEIYRKGSISVFEVDG
KAT8(mouse) GKEIYRKSNISVYEVDG
ESA1(yeast) GNEIYRDDYVSFFEIDG
KAT6B PANEIYRRKDLSVFEVDG
D
KAT6A (8DD5)
KAT5 (2OU2)
KAT8 (6OIO)
KAT7 (5GK9)
KAT8 (mouse, 2Y0M)
ESA1 (yeast, 1FY7)

**Supplementary Figure S1: XL-MS data shows PROTAC-dependent crosslinks** (linked to Figure 1A)

**A-** KAT6A-CRBN inter-crosslinks differentiate between compounds. Heatmap describing the normalized MS intensity of identified crosslinks between KAT6A and CRBN.
**B-** Crosslink abundance correlates with compound potency. Swarm plot displaying the log2 normalized intensity of the identified KAT6B-CRBN crosslink across different compounds, ordered by increasing activity. Individual data points represent independent replicates, and horizontal lines denote the mean. A clear increase in crosslink intensity is observed for more potent compounds.
**C-** Sequence alignment of the KAT catalytic domain. Multiple sequence alignment highlighting the region flanking the target lysine residue. While the surrounding sequence environment is highly conserved across the KAT family, this specific amino acid position exhibits high variability among analogues (e.g., asparagine in KAT6A vs. glycine in KAT6B).
**D-** High tertiary structural conservation of the crosslinked region allows crosslink imputation**.** 3D structural overlay of experimental KAT family structures alongside a Boltz-2 predicted model of KAT6B, with an arrow indicating the spatial position of the target lysine. A magnification of the region is shown in the lower panel. Despite the localized sequence variability shown in (C), the structural backbone of this region is highly conserved across all family members. Spatial equivalence between the experimental structures implies an identical position in both KAT6A and KAT6B, validating the imputation of KAT6B XL-MS constraints to model the active KAT6A ternary complex.

**A**

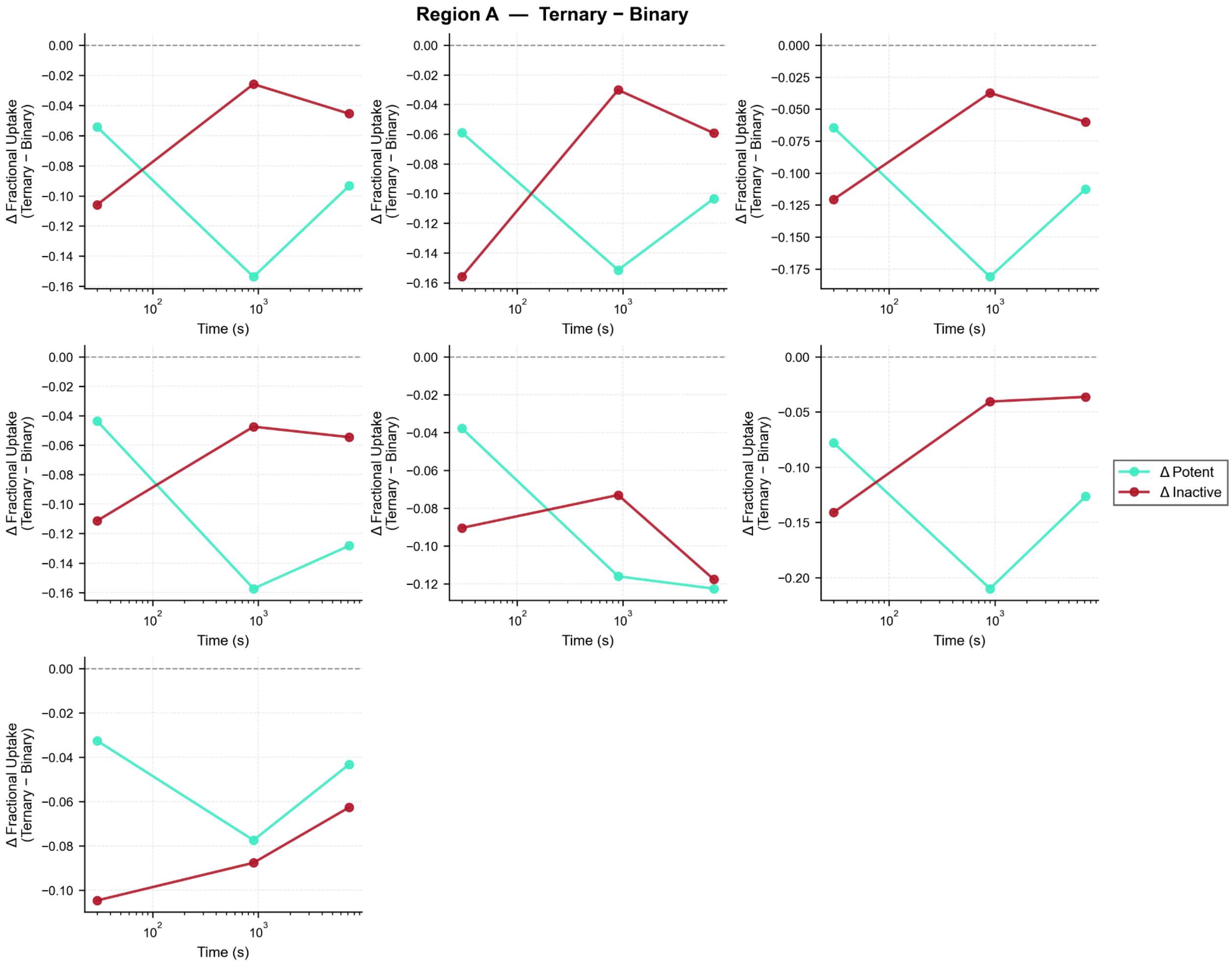

Region A — Ternary − Binary
Δ Fractional Uptake (Ternary − Binary)
Time (s)
Δ Potent
Δ Inactive

**B**

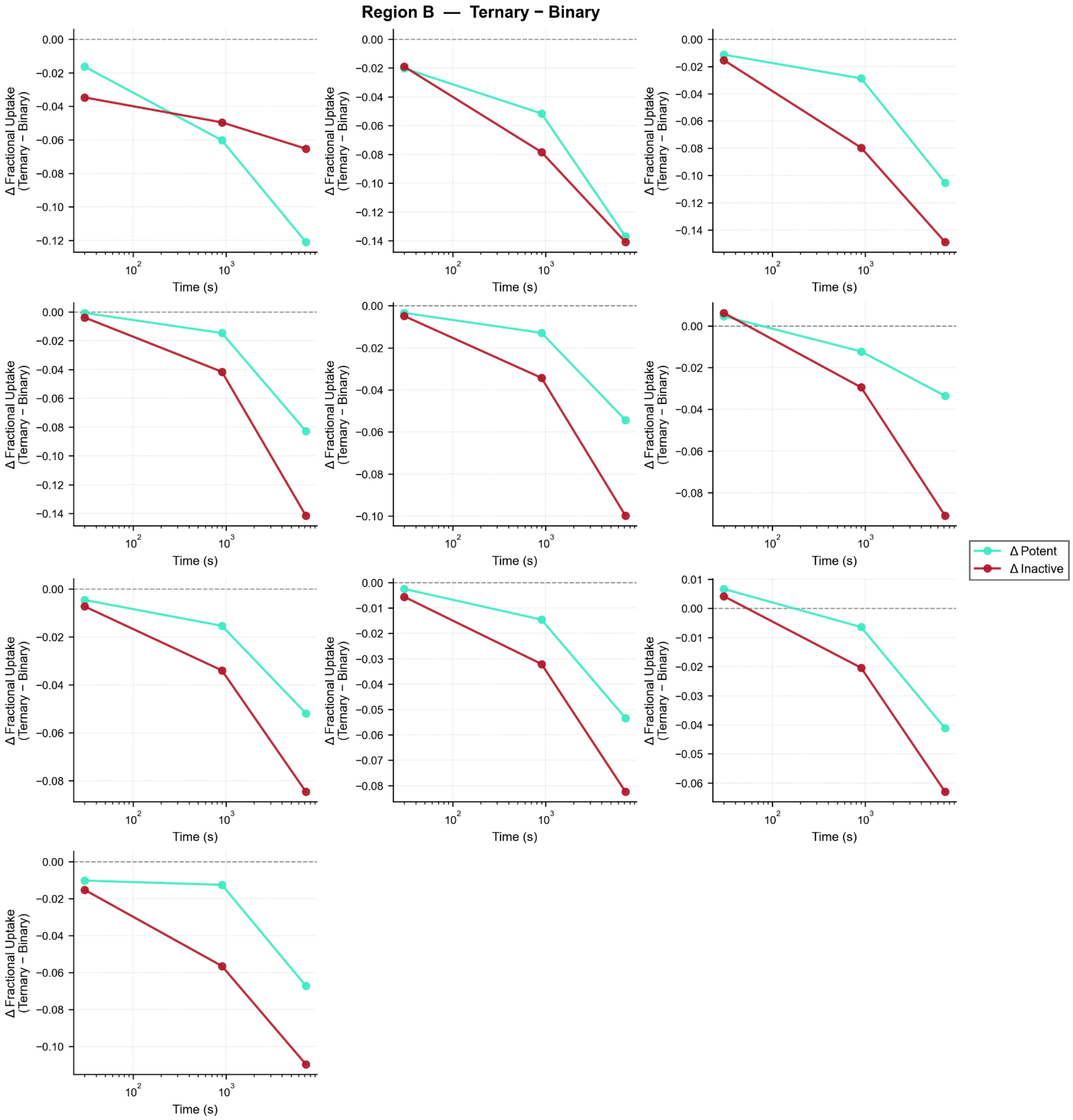


**Supplementary Figure S2: Comparison of deuterium uptake between ternary and binary complexes**

Each graph shows a single peptide and describes the difference in deuterium uptake between ternary (KAT6A-PRTOAC-CRBN) and binary complexes (KAT6A-PROTAC), for either active (green) or inactive (red) compounds. Shown here are only peptides sharing at least 5 amino acids with either Region A (**A**) or Region B (**B**).

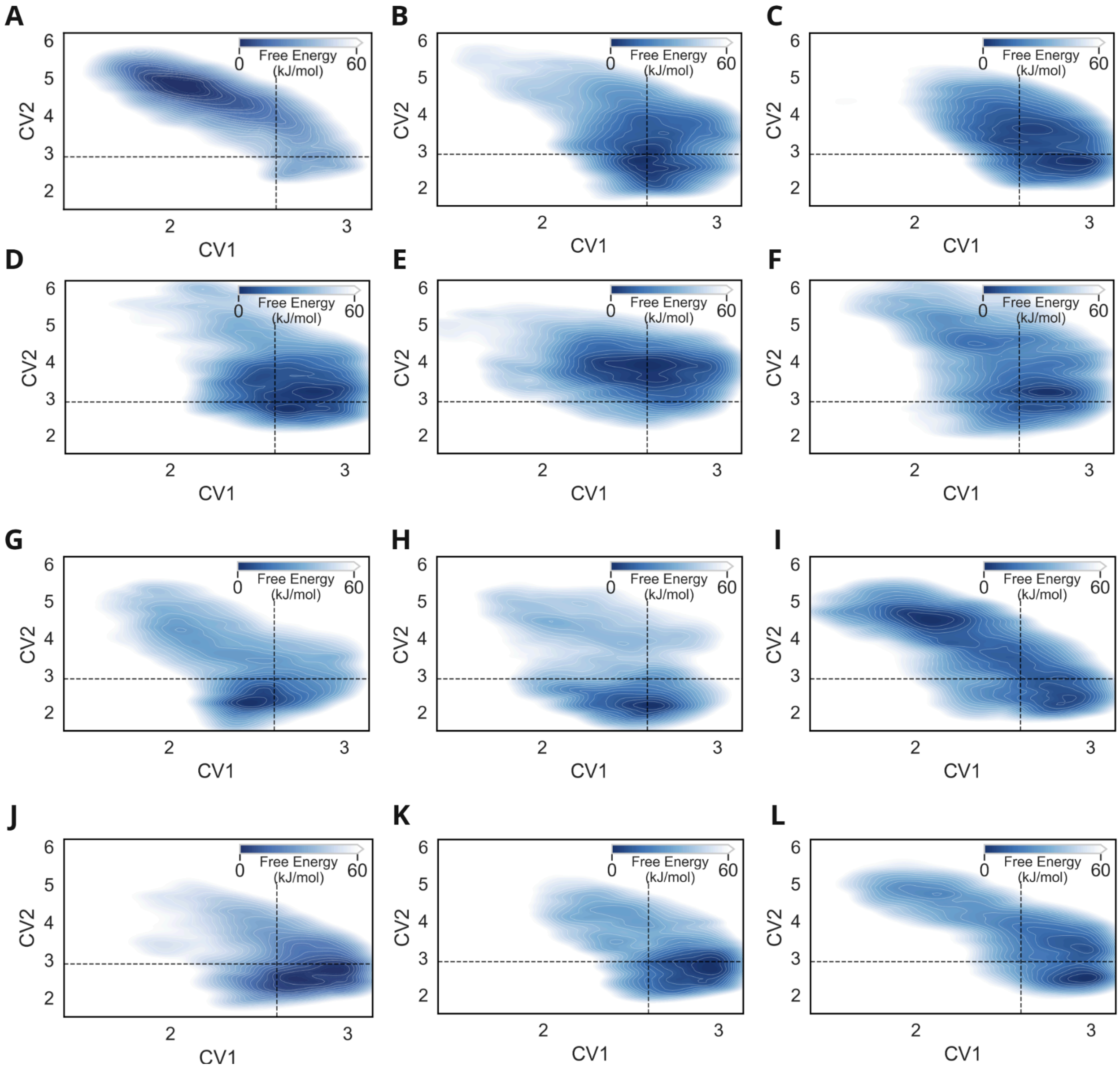


**Supplementary Figure S3: Metadynamics simulations of additional KAT6A degraders** (linked to Figure 2B)

**(A–I)** Estimated free energy surfaces for active compounds and **(J–L)** inactive compounds, plotted as a function of the two collective variables selected for the metadynamics simulations. Energy scales are color-coded in kJ/mol. Deep blue regions represent the lowest energy conformations, illustrating the distinct conformational landscapes accessible to active versus inactive degraders.

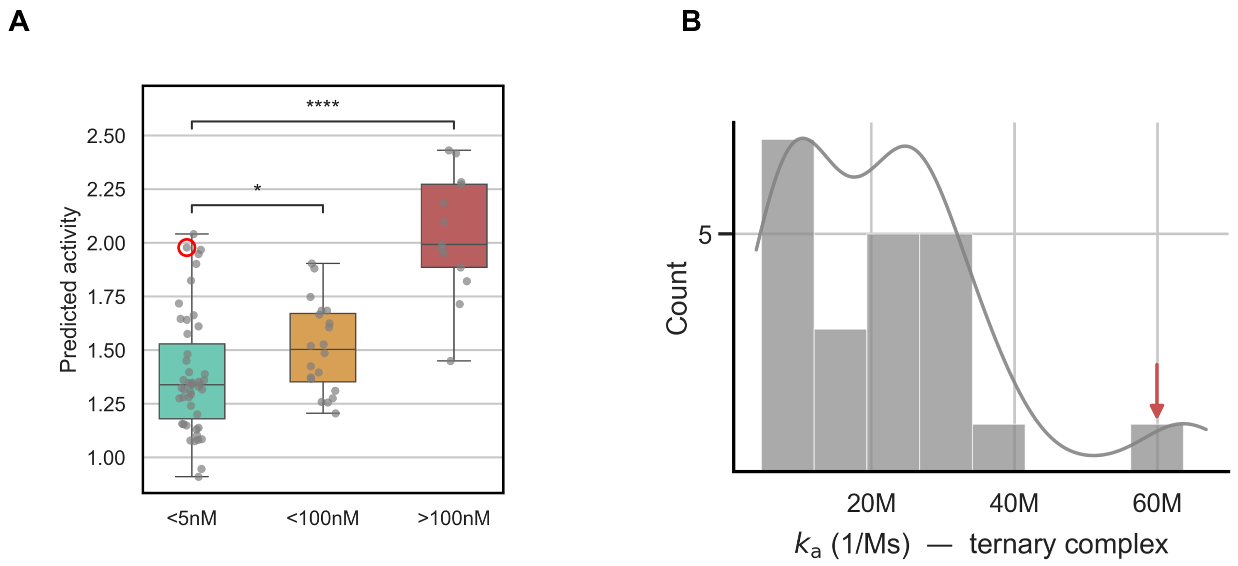


**Supplementary Figure S4: Association rate accounts for AIMS-Rank prediction inaccuracy**

**A-** AIMS-Rank performance summary. The circled data point represents a highly potent compound whose activity was predicted low by the AIMS-Rank model.
**B-** Histogram describing the distribution of association rate ($K_a$) scores measured by SPR. Red arrow points to $K_a$ of the active compound highlighted in S4A.

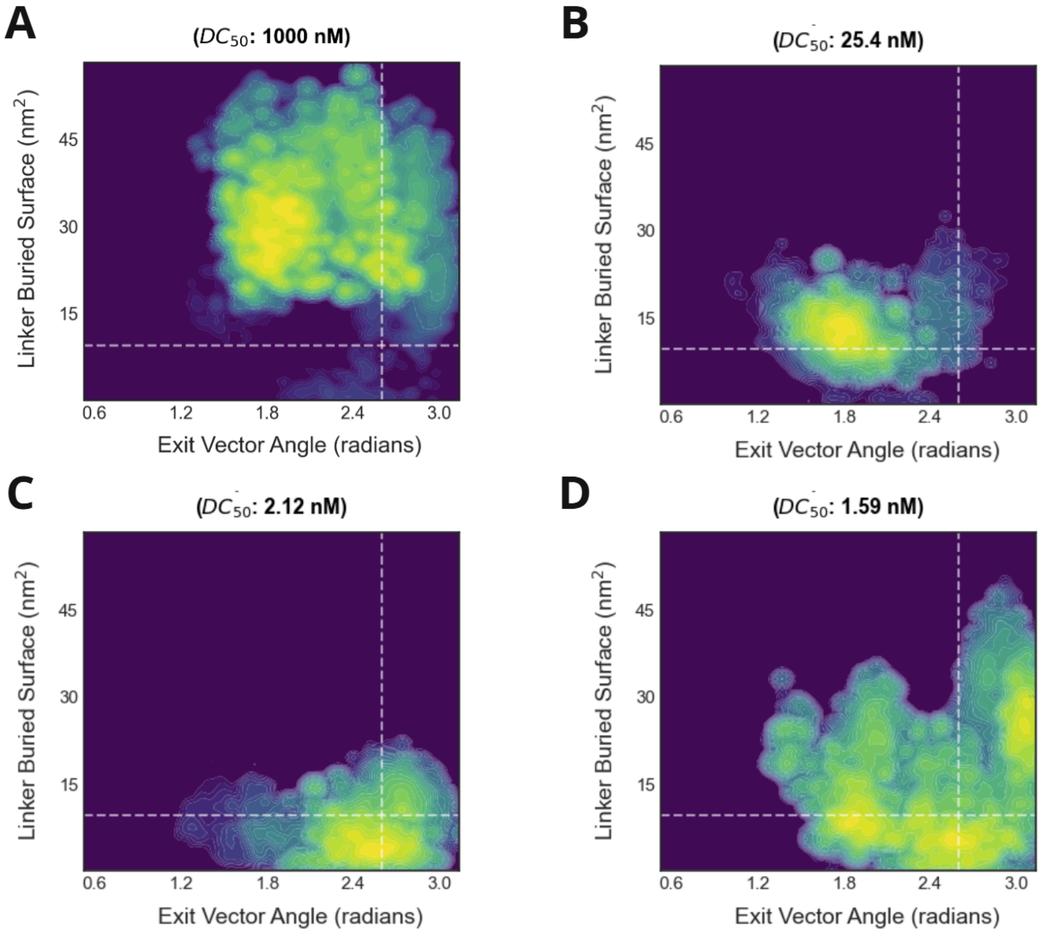


**Supplementary Figure S5: Metadynamics simulations show exit vector angle differentiates between compounds** (linked to Figure 4B)
Estimated free energy landscapes of weak (**A-B**) and strong (**C-D**) degraders, showing larger exit vector angle is associated with stronger degraders.

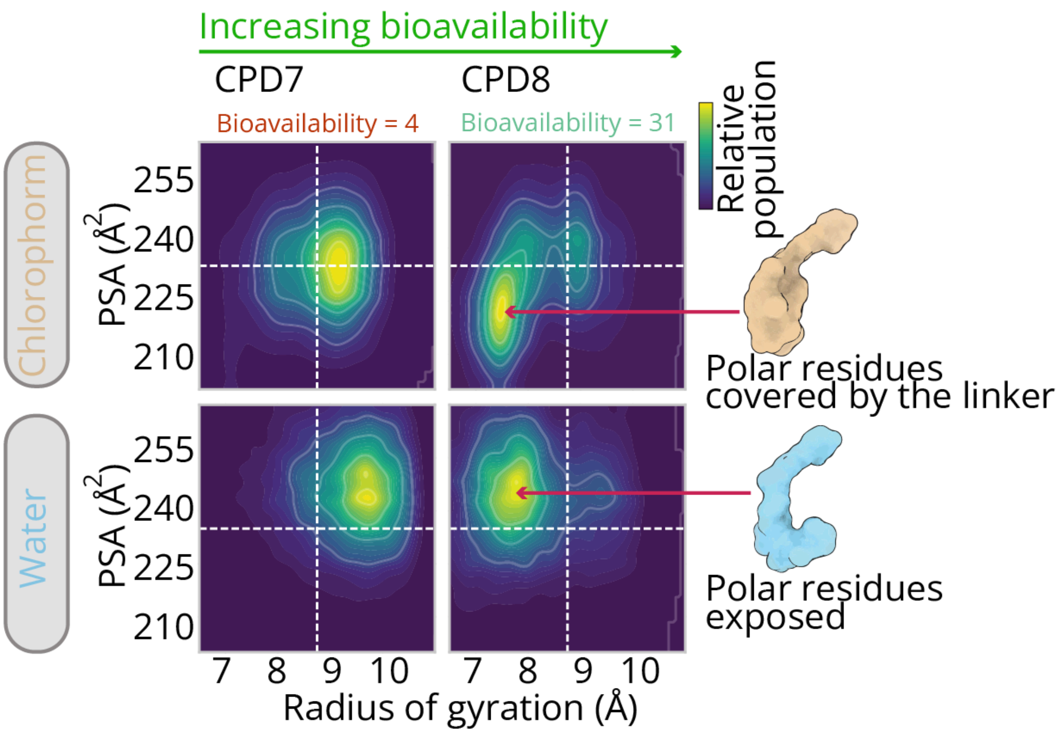


**Supplementary Figure S6: Chameleonic behavior explains differences in compound bioavailability** (linked to Figure 5)
Estimated free energy landscapes of CPD7 (low bioavailability) and CPD8 (high bioavailability) simulations in different environments.

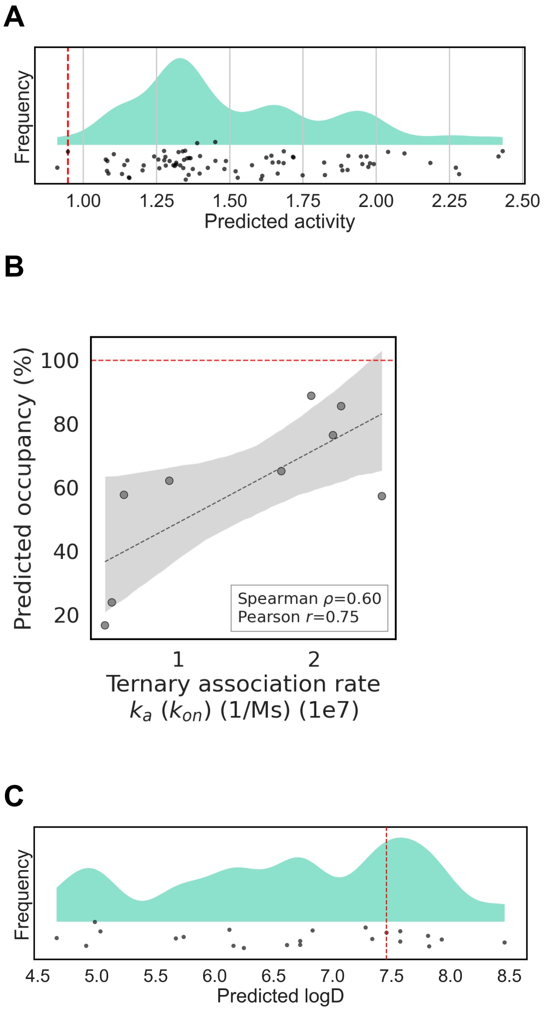


**Supplementary Figure S7: Multi-parameter optimization highlights potential degrader leads**

**A-** Density plot of AIMS-Rank predicted activity, where each point represents a compound and CPD11 is marked in a dashed red line. A lower score indicates better activity.

**B-** CPD11 performance in the complex kinetics model. Same scatter plot as Figure 4C, where the predicted occupancy of CPD11 is marked in a dashed red line (y = 99.96%).

**C-** Density plot of logD prediction, where each point represents a compound and CPD11 is marked in dashed red line. Higher score indicates higher bioavailability.

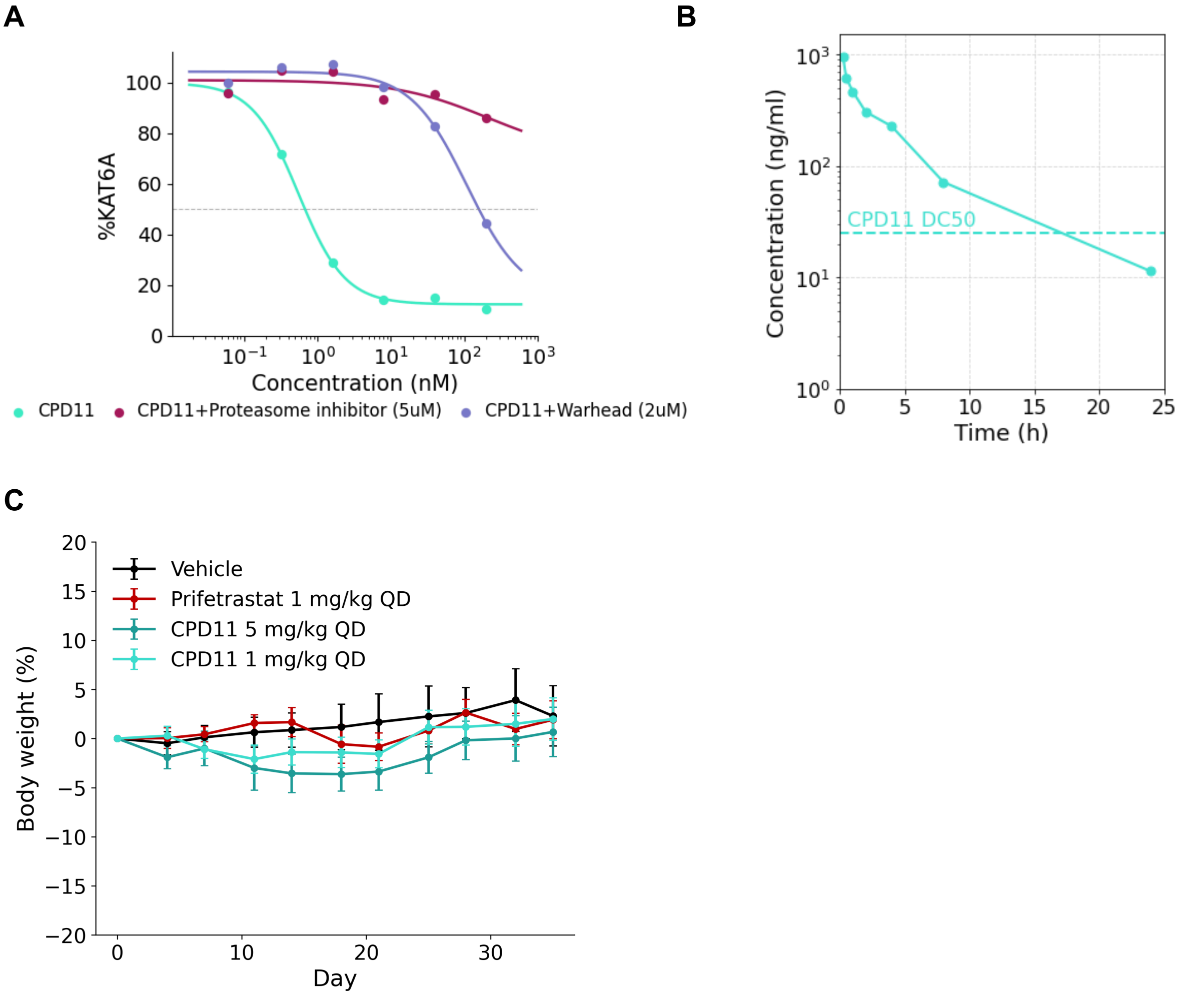


**Supplementary Figure S8**: **CPD11 evaluation *in vitro* and *in vivo*** (linked to Figure 6)

**A-** KAT6A levels were compared across three conditions: treatment of CPD11 alone, CPD11 in combination with the proteasome inhibitor MG132, and CPD11 in combination with an excess of its own KAT6A warhead. KAT6A degradation is dramatically higher with CPD11 treatment alone, validating the correct binding and degradation pathway.

**B-** Plasma pharmacokinetic evaluation of CPD11 in mice (5 mpk, PO). Dashed line represents drug concentration equivalent to in vitro DC50

**C-** No significant body weight loss observed in xenograft mice throughout the treatment period across the different arms.
